# Patch-seq of mouse DRG neurons reveals candidate genes for specific mechanosensory functions

**DOI:** 10.1101/2021.07.07.451447

**Authors:** Thibaud Parpaite, Lucie Brosse, Nina Séjourné, Amandine Laur, Yasmine Mechioukhi, Patrick Delmas, Bertrand Coste

## Abstract

A variety of mechanosensory neurons are involved in touch, proprioception and pain. Many molecular components of the mechanotransduction machinery subserving these sensory modalities remain to be discovered. Here, we combined recordings of mechanosensitive (MS) currents in mechanosensory neurons with single cell RNA sequencing. *In silico* combined analysis with a large-scale dataset enables assigning each transcriptome to DRG genetic clusters. Correlation of current signatures with single-cell transcriptomes provides a one-to-one correspondence between mechanoelectric properties and transcriptomically-defined neuronal populations. Moreover, gene expression differential comparison provides a set of candidate genes for mechanotransduction complexes. *Piezo2* was expectedly found to be enriched in rapidly adapting MS current-expressing neurons, whereas *Tmem120a* and *Tmem150c*, thought to mediate slow-type MS currents, were uniformly expressed in all neuron subtypes, irrespective of their mechano-phenotype. Further knock-down experiments disqualified them as mediating DRG MS currents. This dataset constitutes an open-resource to explore further the cell-type-specific determinants of mechanosensory properties.

## INTRODUCTION

Mechanosensory neurons are involved in multiple sensory modalities, including innocuous tickle, pleasant and discriminative touch, proprioception and kinesthesia as well as various mechanical pain-related sensations such as sharp or deep aching pain and visceral pain. A rich variety of dedicated dorsal root ganglion (DRG) mechanosensory neurons and structured endings innervate our skin, mucosa, muscles, tendons and joints to initiate these mechanosensory sub-modalities (Abraira and Ginty, 2013; Proske and Gandevia, 2012; Roudaut et al., 2012). Physical cues are detected at the sensory nerve endings/auxiliary cells, where specialized mechanosensitive (MS) ion channels convert mechanical stimuli into electrochemical signals (Basbaum et al., 2009; Kefauver et al., 2020).

The discovery from low-throughput patch-clamp screening of PIEZO proteins as genuine force-sensing channels has provided better understanding of mechanical force transduction mechanisms in somatosensory neurons (Coste et al., 2010) as well as in multiple biological systems (Murthy et al., 2017; Wu et al., 2017). PIEZO1 and PIEZO2 form a unique class of a conserved family of nonselective cation channels that are activated by physical stimuli, including pressure, stretch and shear stress (Coste et al., 2012; Ranade et al., 2014a). Both channels generate MS currents with rapid inactivation that convert force into cellular responses on a millisecond time scale. While PIEZO1 is broadly expressed in non-neuronal tissues exposed to pressure and fluid flow, e.g., kidneys, bladder, endothelial cells, and blood cells, PIEZO2 is predominantly expressed in sensory tissues, such as DRG, trigeminal ganglia and Merkel cells (Murthy et al., 2017; Wu et al., 2017). Studies from knock-out (KO) mouse models have shown that *Piezo2* plays a crucial role in innocuous touch sensation (Ranade et al., 2014b) and proprioception (Florez-Paz et al., 2016; Woo et al., 2015). *Piezo2* also plays a more marginal role in mechanical nociception (Murthy et al., 2018b; Szczot et al., 2018). These functions appeared to be conserved in humans, as patients with loss-of-function variants in *PIEZO2* show deficits in touch discrimination and joint proprioception, and failed to develop mechanical allodynia after skin inflammation (Chesler et al., 2016; Szczot et al., 2018).

DRG neurons express a large repertoire of excitatory MS currents classified as rapidly-adapting (RA), intermediately-adapting (IA), and slowly/ultra slowly-adapting (SA/ultra SA) currents, according to their inactivation kinetics to sustained mechanical stimulation (Coste et al., 2010; Drew et al., 2004; Drew et al., 2002; Francois et al., 2015; Hao and Delmas, 2010; Hu and Lewin, 2006; Ranade et al., 2014b; Rugiero et al., 2010). It is now well established that PIEZO2 channels sustain RA MS currents (Coste et al., 2010; Ranade et al., 2014b). Importantly, IA and slow-type MS currents are largely unaffected in *Piezo2* KO DRG neurons (Ranade et al., 2014b), demonstrating that as-yet-unknown MS channels must account for the other MS currents. It is worthwhile to identify these channels since many sensations related to innocuous and noxious mechanical stimuli are independent from *PIEZO2* in humans (Case et al., 2021; Chesler et al., 2016; Szczot et al., 2018). Recently, two other genes, namely *Tmem120a* (Tacan) and *Tmem150c* (Tentonin 3), have been proposed to encode ion channels sustaining slow MS currents in DRG neurons (Beaulieu-Laroche et al., 2020; Hong et al., 2016), but some of these findings are controversial (Anderson et al., 2018; Dubin et al., 2017). Hence, despite significant advances, the full panel of MS channels involved in mechanosensation has yet to be established.

Identification of MS channels is hampered by the large heterogeneity of DRG sensory neurons. This heterogeneity has been well documented by large scale single-cell transcriptomics enabling gene expression-based classification (Li et al., 2016; Sharma et al., 2020; Usoskin et al., 2015; Zeisel et al., 2018). These studies have identified eight main classes of sensory neurons, including low-threshold mechanoreceptors, proprioceptors, thermoreceptors, nociceptors, itch sensitive neurons and type C low-threshold mechanoreceptors, which can be further subdivided into up to 17 clusters identified genetically (Zeisel et al., 2018). A limitation of large-scale classification based on unsupervised grouping of neurons with similar expression profiles is the lack of direct functional correlates. Indeed, this would require a prior functional characterization of individual neurons by patch-clamp experiments, a laborious step incompatible with large samples of neurons. Nevertheless, the direct correlation of single cell transcriptomes with functional mechanosensory properties could serve as a basis for identifying the genes involved in mechanotransduction.

Here, we combined recordings of MS currents in cultured mouse DRG neurons with single cell RNA sequencing. *In silico* analysis of collected data with a previous large-scale dataset allowed to link MS current subtypes to molecular neuronal clusters. These data made it possible to precise the distribution of the four different types of MS currents in molecularly defined populations of DRG neurons. Moreover, gene expression differential comparison provided a list of candidate genes for mechanotransduction complexes. This dataset combining patch-clamp electrophysiology and quantitative single cell RNA sequencing constitutes an open-resource to explore further the cell-type-specific determinants of mechanosensory properties.

## RESULTS

### Coupling patch-clamp recording of MS currents with single cell RNA sequencing

To identify molecular features associated with specific MS phenotypes, we combined single cell RNA-sequencing (scRNAseq) profiling with electrophysiological characterization of MS currents in individual DRG neurons. A representation of the methodological procedure used for data collection is depicted in figure 1A. Briefly, sensory neurons from mouse DRG cultures were recorded using the whole-cell patch-clamp mode in order to characterize the properties of excitatory MS currents in response to cell poking with a mechanical probe (Coste et al., 2007; Coste et al., 2010; Drew et al., 2002; Hao and Delmas, 2010; Hu and Lewin, 2006; McCarter et al., 1999; Michel et al., 2020; Rugiero et al., 2010). After functional characterization, the content of each cell was harvested through the recording pipette prior to the generation of cDNA libraries and RNA sequencing (see M&M for detailed procedure). Using this approach, we selected 62 DRG neurons covering the main types of MS currents (Fig.1B) for cDNA library preparation, of which 53 cells proved suitable for scRNAseq analysis. A total of 1.6 billion paired-end reads were generated, among which 1.4 billion were correctly aligned to mouse genome, giving an average of 25.9 million mapped reads for a given neuron. The mean number of detectable genes per cell was 13,359 ± 1,478. Analysis of reference genes showed no differences between samples, which attested the absence of batch-effect between the sequencing runs (Fig.S1A). Quality metrics values from sequenced neurons were within the same range than large scale DRG scRNAseq studies (Li et al., 2016; Usoskin et al., 2015; Zeisel et al., 2018), attesting that patch-clamping/poking the neurons prior to RNA sequencing analysis did not impact gene detection (Fig.S1B). Overall, the depth of our molecular analysis allowed us to quantitatively assay the expression of genes in each DRG neuron and to compare expression differences of any of these genes in light with mechanical properties.

**Figure 1.**
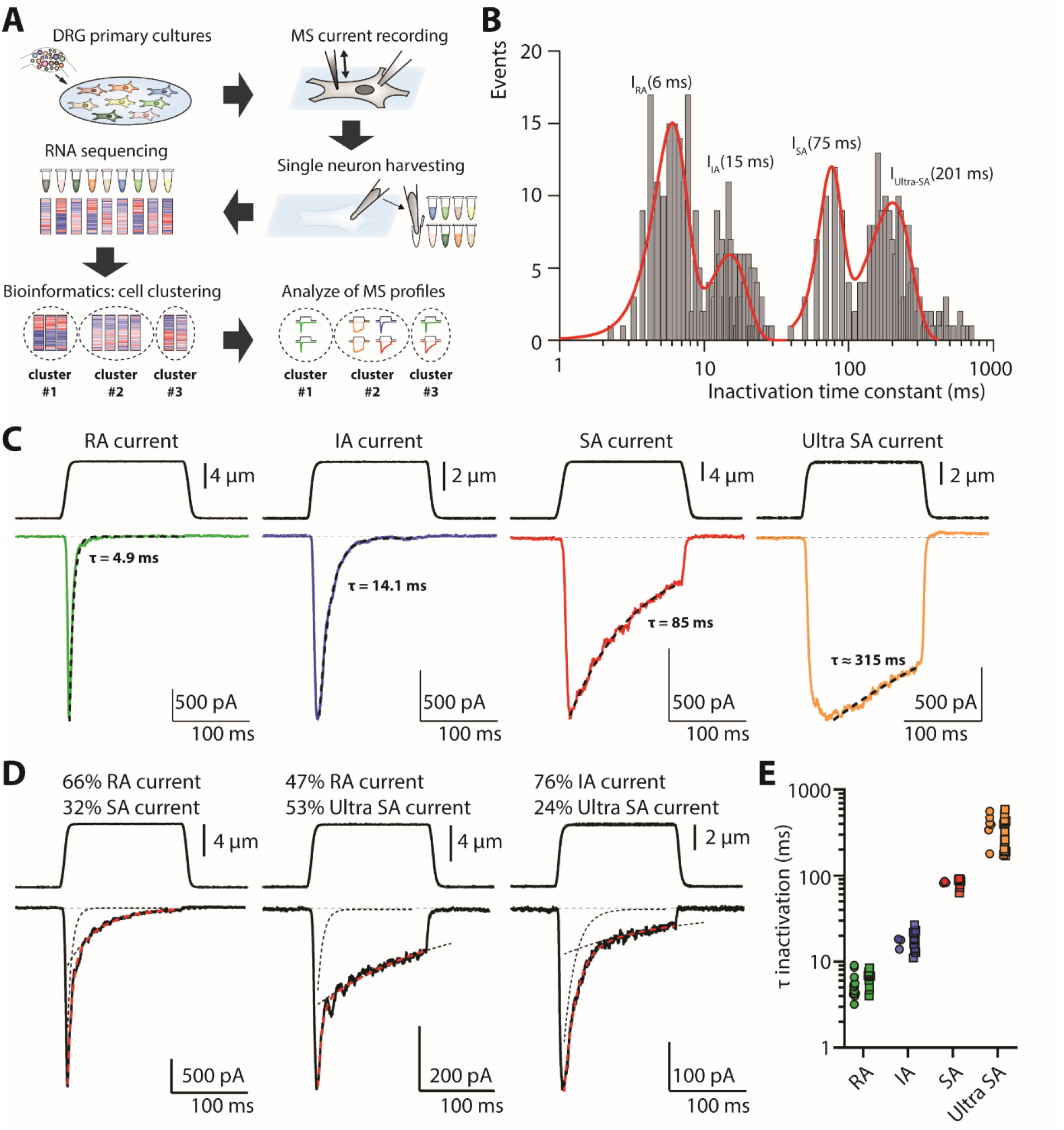
Single cell RNAseq coupled to MS current characterization. (**A**) Schematic representation of the methodology used in this study. Fifty-three individual neurons were successfully sequenced following this procedure. (**B**) Frequency distribution of the inactivation time constants determined from mono- and bi-exponential fits of MS currents. Data collected over 312 DRG neurons, voltage clamped at -80 mV and stimulated with a mechanical probe. Gaussian distribution identified four centers of inactivation time constants. Bin width was 0.5 ms for I_RA_ and I_IA_, and 10 ms for I_SA_ and I_Ultra-SA_. (**C**) Representative examples of the distinct types of MS currents with inactivation time-course fitted with a mono-exponential function (black dashed line). (**D**) Examples of MS currents fitted with bi-exponential functions (red dashed line). Black dashed lines show each mono-exponential component extracted from the fit. For **C** and **D**, upper traces represent the mechanical probe displacement and lower traces the recorded currents at a holding potential of -80 mV. (**E**) Inactivation time-constant (τ) for all identified current components in scRNAseq neurons with monophasic (circle) or biphasic (square) MS currents.

### Categorization of MS current-expressing neurons prior to transcriptome profiling

To tentatively classify mechanosensory neurons based on their mechanical properties, we examined the kinetics of MS currents over a large population of mouse DRG neurons (n = 312), not intended for scRNAseq. Previous studies have identified 3 to 4 kinetically distinct MS currents in rat and mouse DRG neurons, which may reflect the activation of non-uniform populations of ion channels (Coste et al., 2010; Drew et al., 2004; Drew et al., 2002; Francois et al., 2015; Hao and Delmas, 2010; Hu and Lewin, 2006; Ranade et al., 2014b; Rugiero et al., 2010). Based on exponential fit of inactivation kinetics, we identified four kinetically distinct MS currents with gaussian mean value centered at 5.9 ± 1.6, 15.1 ± 4.5, 75.4 ± 13.8 and 200.9 ± 66.8 ms (mean ± SD) (Fig.1B). These currents were referred hereafter as rapidly adapting (RA; τ_inac_ ≤ 10 ms), intermediately adapting (IA; 10 ms < τ_inac_ ≤ 30 ms), slowly adapting (SA; 30 ms < τ_inac_ ≤ 110 ms) and ultra-SA (τ_inac_ > 110 ms). Prototypical recordings of these currents are depicted in figure 1C. It is worth noting that a substantial part of mechanosensory DRG neurons exhibited more than one single current component, potentially reflecting the activation of two or more populations of distinct MS channels. For analytical purpose, we only considered mixed currents best fitted by bi-exponential functions. The contribution of each component to the whole MS current was estimated from the amplitude of the respective exponential fit (Fig.S2, see M&M for detailed procedure). Representative examples of biphasic currents classified as RA/SA, RA/ultra-SA and IA/ultra-SA are illustrated in figure 1D.

This kinetics-based categorization made it possible to classify our successfully sequenced DRG neurons. Thus, from our sample of 53 scRNAseq neurons, 49% displayed a monophasic MS current, while the remaining 45.5% showed a mixed current, with variable MS current components. Only 3 scRNAseq neurons were unresponsive to mechanical stimulus, but this does not reflect physiological proportion (Coste et al., 2010; Hu and Lewin, 2006). Average values of τ_inac_ were 5.9 ± 0.3, 18.8 ± 1.4, 82.2 ± 2.8 and 347.7 ± 26.5 ms for RA, IA, SA and ultra-SA current, respectively (n = 26, 15, 10 and 24; mean ± sem) (Fig.1E). The detailed profiles of MS currents in individual neurons are described in Table S1.

### Mapping Patch-seq transcriptomes of MS neurons to single-cell RNA-seq datasets

Previous scRNAseq studies performed on large samples of neurons have used gene expression profile analysis to sort out DRG neurons into molecularly defined neuronal subclasses (Li et al., 2016; Sharma et al., 2020; Usoskin et al., 2015; Zeisel et al., 2018). To achieve high-quality classification of our neuron sample, we merged our data with the dataset of 1580 scRNAseq DRG neurons from Zeisel and collaborators, which is available as open source (Zeisel et al., 2018). Then, we performed a joint analysis using mutual nearest neighbors (MNN) correction by focusing on the expression magnitude of the 211 most highly enriched genes across DRG neuron clusters (Zeisel et al., 2018) (Fig.S1C). This procedure allowed to classify each neuron from our sample in genetic clusters identified previously (Zeisel et al., 2018) (Table S1). The merged datasets were mapped using t-distributed stochastic neighbor embedding (tSNE) projection (Fig.2A). Neurons from our sample spread across seven out of eight functionally distinct DRG neuron populations, some of which are composed of multiple genetic clusters whose functional specificity remains to be determined. Eight neurons were classified into the NF group of neurons typically considered as low-threshold mechanoreceptors (LTMR) and proprioceptors. Six neurons belong to NP1 population related to polymodal nociceptors, whereas eleven neurons were assigned to NP2 population and a single neuron to NP3 population, which are related to itch-specific sensory neurons (Cavanaugh et al., 2009; Dong and Dong, 2018; Han et al., 2013; Wang and Zylka, 2009). Nine neurons were assigned to PEP1, a population of C-type thermo-nociceptors, and sixteen neurons to PEP2, a population of lightly myelinated Aδ nociceptors (Usoskin et al., 2015). Two neurons segregated into TH neuron population, representing C-low threshold mechanoreceptors (C-LTMR), which contribute to pleasant touch (Li et al., 2011; Olausson et al., 2010) and mechanical allodynia (Seal et al., 2009). Finally, none of our scRNAseq neurons belong to TrpM8 population. These neurons represent a labeled line for cold sensation, and their genetic ablation in mice has no effect on mechanical behaviors (Knowlton et al., 2013).

**Figure 2.**
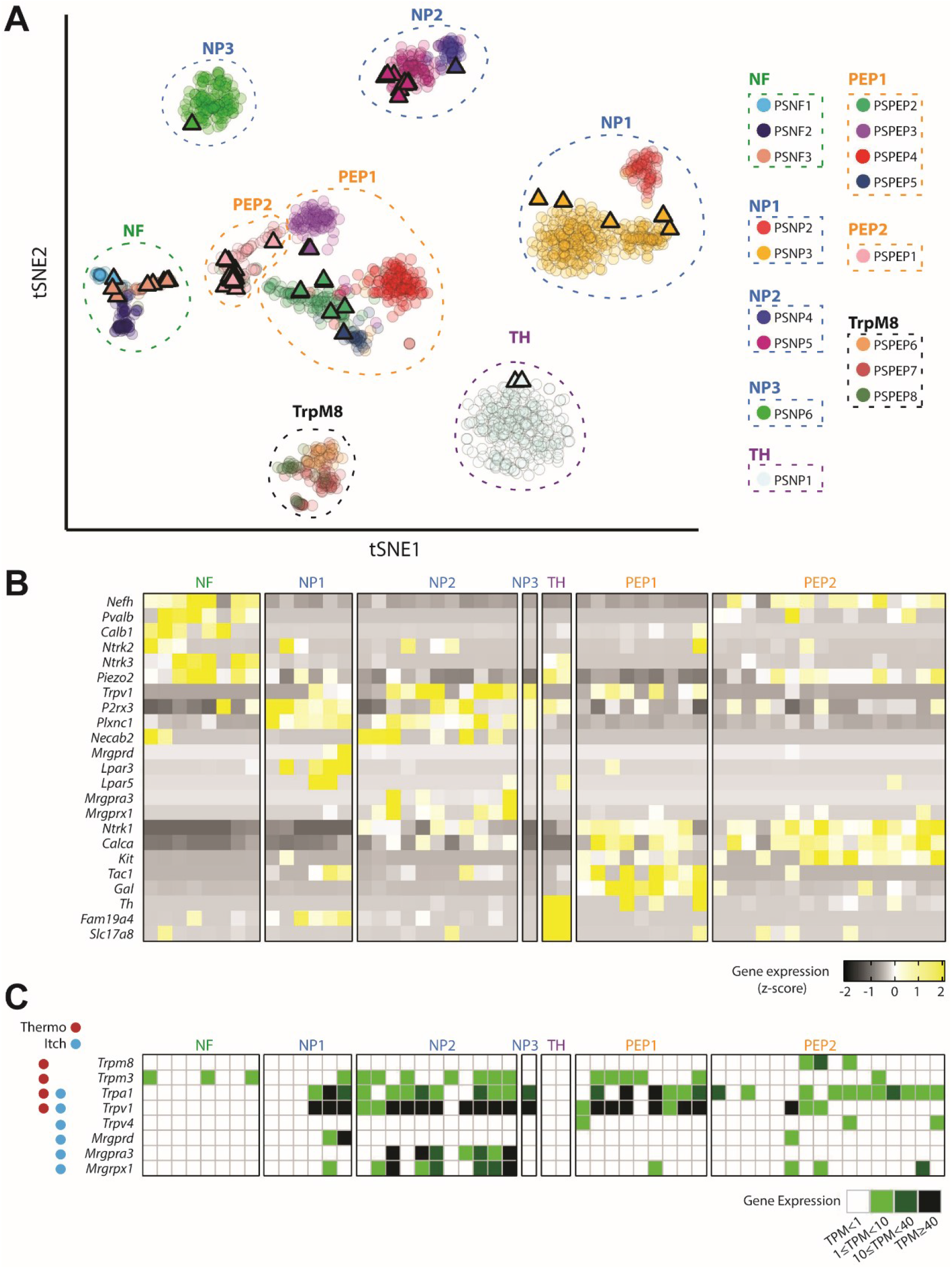
Cluster analysis of scRNAseq neurons. (**A**) Visualization using t-SNE embedding of all cells colored by cluster identity. Triangles represent the 53 scRNAseq neurons of this study and circles the 1580 neurons from the study of Zeisel and collaborators. Main populations of neurons are surrounded by dashed lines. See **Fig.S1C** for the heat map of the expression of genes used for joint analysis. (**B**) Heat map of expression of marker genes in neurons grouped by population. (**C**) Expression level of genes coding for thermosensitive and pruriceptive molecular sensors in neurons grouped by population. NF: Neurofilament; NP: Non-peptidergic; TH: Tyrosine hydroxylase; PEP: Peptidergic.

Expression of marker genes and genes coding for thermosensitive and pruriceptive molecular sensors confirmed the appropriate mapping of scRNAseq neurons from our sample (Fig.2B-C). NF LTMR and proprioceptors express the specific markers *Nefh* (neurofilament heavy chain), *Pvalb* (parvalbumin), *Calb1* (calbindin1), *Ntrk2* and *Ntrk3* (TrkB and C), but thermosensitive and pruriceptive itch molecular sensors were not detected. NP1-3 sensory neurons express *P2rx3* (P2X3), *PlnxC1* (plexin C1) and *Necab2* (N-terminal EF-hand calcium binding protein 2). The itch related genes *Lpar3* and *Lpar5* (lysophosphatidic acid receptors 3 and 5), and *Mrgprd* (MGRPRD) are expressed in NP1 neurons, whereas NP2 population expresses *Mrgpra3* and *Mrgprx1* (MRGPRA3 and X1). PEP neurons express *Calca* (CGRP) and *Kit* (c-Kit) but differ in the expression of *Gal* (galanin prepropeptide) for PEP1 C-type thermo-nociceptors and *Nefh* for PEP2 lightly myelinated Aδ nociceptors. The heat noxious sensors *Trpa1, Trpv1* and *Trpm3* (Vandewauw et al., 2018) were strongly detected in NP1, NP2 and PEP1 C-type neurons compared with PEP2 Aδ-type neurons. This is consistent with previous observation in skin-nerve recording studies that showed a higher proportion of C-type than Aδ-type fibers that respond to noxious heat (Cain et al., 2001; Koltzenburg et al., 1997). Finally, the TH C-LTMR characterized by expression of *Th* (tyrosine hydroxylase), *Fam19a4* (TAFA4) and *Slc17a8* (Vglut3), are devoid of thermosensitive or itch-related molecular sensors.

### Distribution of MS currents in genetically defined DRG subclasses

The characterization of MS current types expressed in scRNAseq neurons allowed us to examine their distribution in genetically defined DRG populations (Fig.3A-B). Except for NF neurons, which exhibited almost exclusively the RA current, all other molecularly defined DRG subclasses displayed more than one type of MS current. However, the prevalence of each current component differed among DRG groups. The Figure 3C shows the proportion of each MS current type in genetically defined DRG neuron populations. The polymodal nociceptor NP1 subclass displayed all four types of MS currents, with individual contributions ranging from 10 to 33%. The itch specific NP2 subclass expressed almost no SA current and showed a higher amount of RA current (42%) compared with IA and ultra-SA currents (35 and 20%, respectively). The C-type thermo-nociceptor PEP1 subclass had prominent IA current (56%), a lower amount of RA current (33%), with no or little SA and ultra-SA current components, respectively. The Aδ nociceptor PEP2 subclass almost lacked IA current, and expressed prominent ultra-SA current (39%) and robust RA and SA currents (34 and 23%, respectively). Finally, C-LTMR TH neurons showed both RA and ultra-SA currents. Although our small sample size precluded definite conclusion about the nature of MS current in C-LTMR TH neurons, our data are consistent with previous work indicating that TH neurons mainly express RA and ultra-SA currents, or a mixture of both (Delfini et al., 2013). Altogether, these results revealed that the genetically defined DRG subclasses have different complements of MS currents, which may reflect some heterogeneity and functional differences within a given DRG subclass.

**Figure 3.**
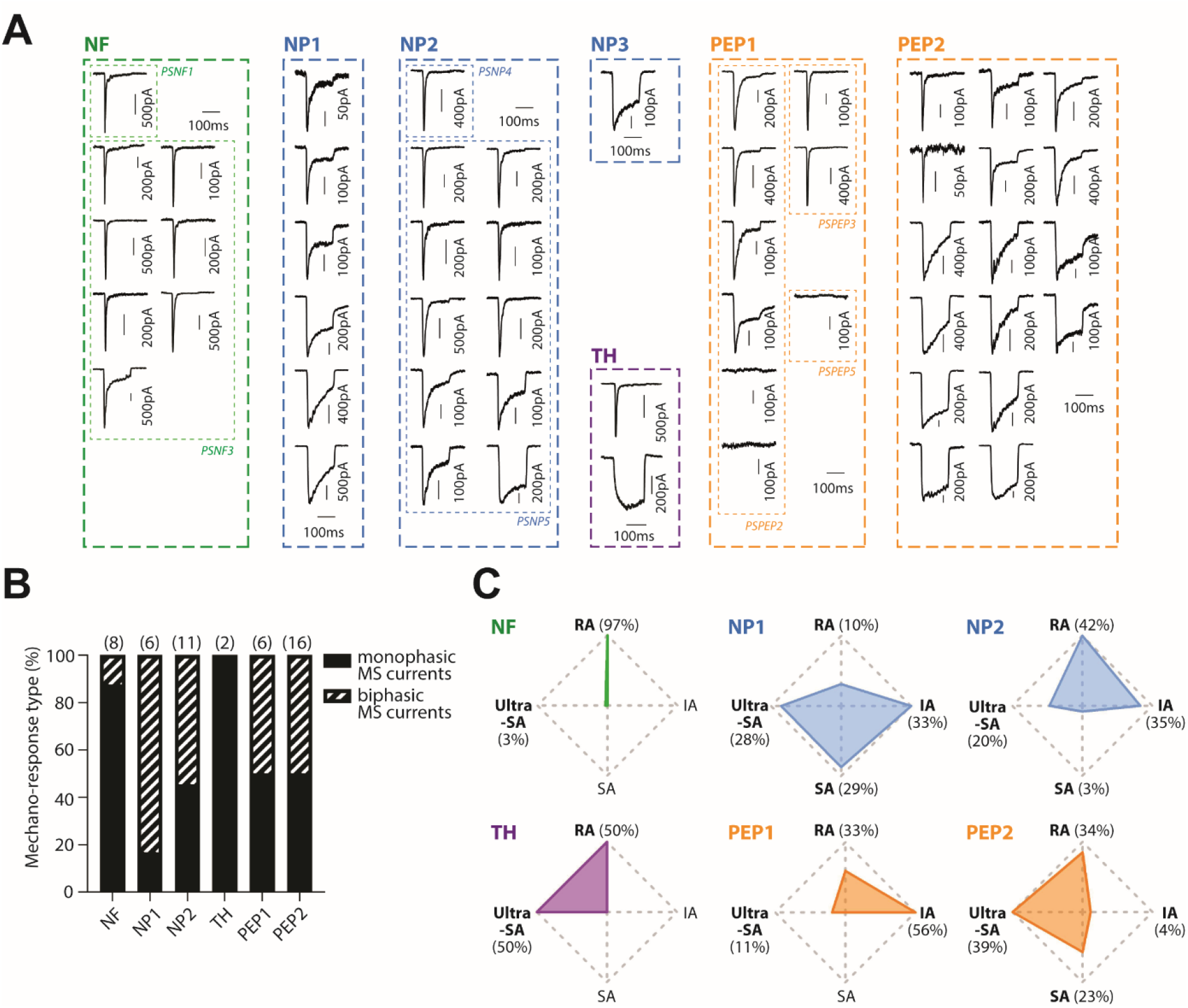
Distribution of MS currents among genetic clusters of DRG neurons. (**A**) Traces of MS currents recorded in the 53 scRNAseq neurons and grouped by neuronal genetic population. Mechanical stimulation traces are not shown for clarity sake. (**B**) Proportion of MS neurons expressing monophasic and biphasic MS currents in each genetic population. Number of neurons for each population is in brackets. (**C**) Relative contribution of MS current types to the total MS current in each genetic population. Numbers of neurons are as in **B**.

### Expression analysis of genes hypothesized to encode MS channels

It has been remarkably difficult to identify candidate excitatory MS channels in mammals. With the notable exception of PIEZOs and of TMCs in auditory hair cells (Corey et al., 2019), other putative candidates have not been confirmed or might be modulators of mechanosensation rather than being directly involved in force transduction. A prerequisite for a candidate MS channel gene is to be found expressed in neurons displaying the purposed MS current. Therefore, we examined whether the expression of genes that have been proposed to encode excitatory MS channels in mammals correlates with the presence of a given MS current subtype across the scRNAseq neurons (Fig.4A). As expected, *Piezo2* was detected in all neurons exhibiting the RA current. However, *Piezo2* was also detected in neurons for which no RA current could be extracted. *Tmem120a*, which has been proposed to sustain MS currents with slow kinetics in DRG neurons (Beaulieu-Laroche et al., 2020) was detected at high level in all neuron types, including neurons lacking MS current. *Tmem150c* (Tentonin 3), which has been proposed to encode MS channels with slow kinetics in DRG and nodose ganglion neurons (Hong et al., 2016; Lu et al., 2020), was detected in most neurons, regardless of their mechanical signature. *Tmem87a*, a gene involved in melanoma adhesion/migration and sufficient to reconstitute MS currents in heterologous systems (Patkunarajah et al., 2020), was detected in some neurons, irrespective of the MS current subtypes. All three *Tmem63* genes which belong to an evolutionary conserved family of MS channel coding genes across eukaryotes (Murthy et al., 2018a) were differentially expressed. Tmem63a was detected in few neurons but *Tmem63b* and *Tmem63c* were found in most MS current-expressing neurons. Finally, *Piezo1* was sporadically detected, while *Tmc1* and *Tmc2*, which contribute to the pore of MS channels in auditory and vestibular hair cells (Jia et al., 2020; Pan et al., 2018; Pan et al., 2013) were almost not expressed. Together, these results lend credence to the idea that PIEZO1 and TMC1 and 2 are unlikely to contribute to MS currents in DRG neurons. They also indicate that the other candidate genes are expressed in different MS current neuron families with no apparent clustering/specific expression.

### Enrichment of transcripts as a mean to identify MS channel gene candidates

The above data failed to reveal specific expression of a given MS channel gene in a particular neuronal group. Therefore, we analyzed the differential gene expression to assess whether some transcripts showed statistically over-representation (e.g. enrichment) in neurons clustered by current types. The plots illustrated in figure 4B provide graphical views of the enrichment of each MS channel gene in neurons with RA, IA, SA and ultra-SA components. *Piezo2* is differentially expressed (*P = 0*.*015*) in neurons expressing RA currents, exhibiting a 2.3-fold enrichment. This result indicates that relative enrichment rather than specific expression may be a pertinent criterion when selecting putative MS current genes. Remarkably, none of the other MS channel gene candidates were found to be differentially expressed (*P > 0*.*05*) in IA, SA or ultra-SA MS current-expressing neurons. This lack of enrichment raises doubt about their possible contribution to specific MS currents in DRG neurons. As this result is intriguing regarding the recent literature, we set up proof-of-principle experiments to probe the functional relevance of *Tmem120a* and *Tmem150c* in MS currents.

**Figure 4.**
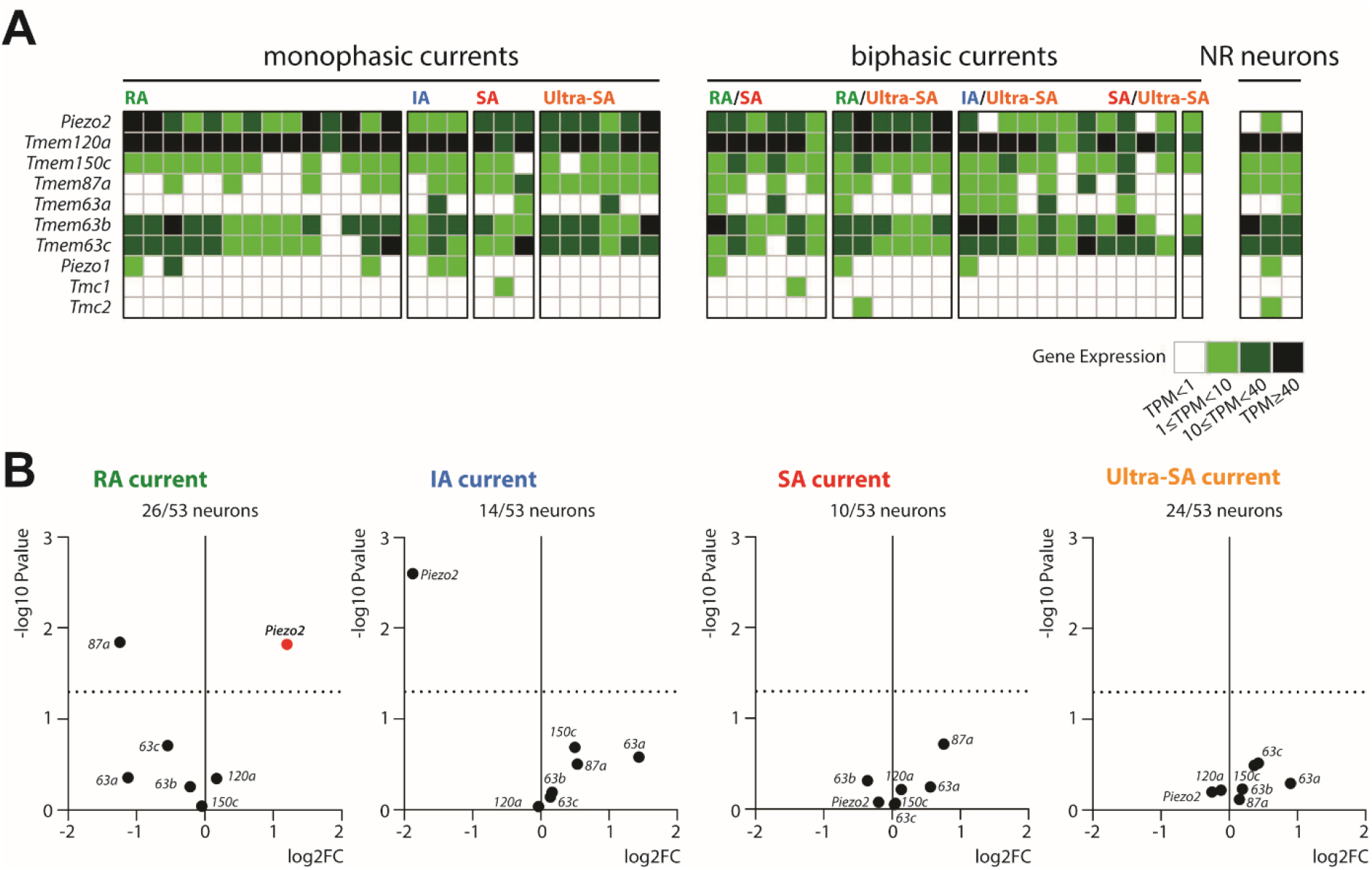
Expression analysis of genes proposed to encode MS channels. (**A**) Expression level of genes related to MS channels in scRNAseq neurons grouped by MS current type, as indicated. NR, non-responsive to mechanical stimulation. (**B**) Differential expression plots of genes related to MS channels in neurons grouped by MS current type, as specified. Dashed line represents p value of 0.05. For *Tmem* genes, only the label after *Tmem* is indicated for clarity.

### Inhibition of *Tmem120a* expression does not alter DRG neuron MS currents

Our scRNAseq data indicated that *Tmem120a* is broadly expressed in all neuron subclasses. We further characterized *Tmem120a* expression in DRG slices using the RNAscope *in situ* hybridization (ISH) technology coupled with immunostaining for the broader nociceptive and non-nociceptive markers, Peripherin and NF200, respectively (Goldstein et al., 1991; Padilla et al., 2007) (Fig.5A-C, Fig.S3). *Tmem120a* transcripts were detected in 75.0 % of NF200-positive neurons and 67.4 % of peripherin-positive DRG neurons (Fig.5B). Moreover, analysis of the cross-sectional area distribution revealed that *Tmem120a* mRNA was detected in the majority of DRG neurons over a large range of cell soma diameters (Fig.5C). Altogether, these results confirm that *Tmem120a* is widely expressed in the different subpopulations of sensory neurons.

**Figure 5.**
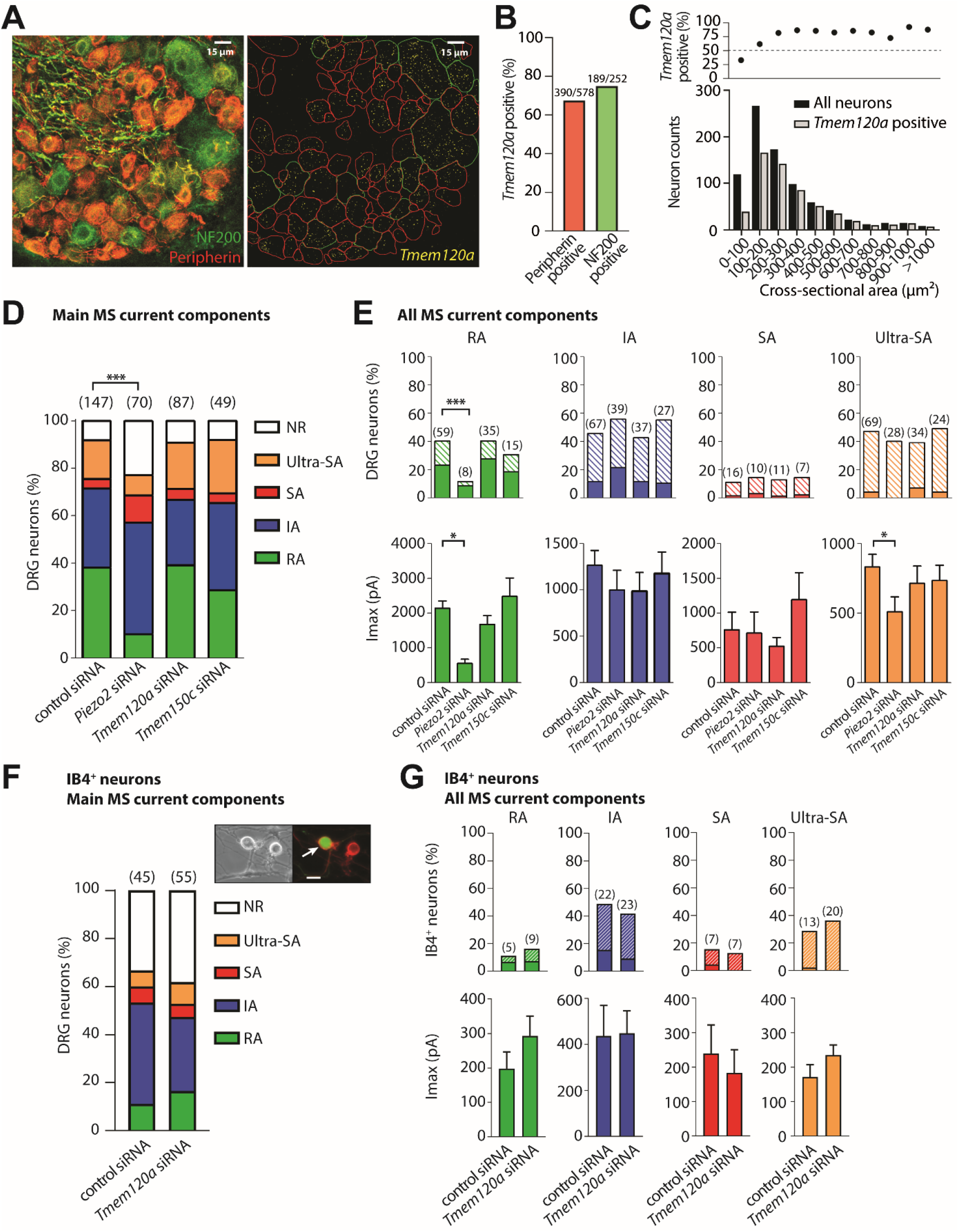
*Tmem120a* and *Tmem150c* do not contribute to DRG MS currents. (**A**) Representative images of fluorescent *in situ* hybridization for *Tmem120a* (right panel) in DRG neurons immuno-stained for peripherin and NF200 (left panel). (**B**) Percentage of DRG neurons expressing *Tmem120a* mRNA in Peripherin- or NF200-positive populations. (**C**) Cross-sectional area distribution of *Tmem120a* mRNA-positive neurons. The top panel shows the percentage of *Tmem120a* positive neurons. **(D)** Quantification of the prevalence of RA, IA, SA and ultra-SA currents among DRG neurons electroporated with control, *Piezo2, Tmem120a* or *Tmem150c* siRNAs. Neurons responding with biphasic MS currents were classified according to the MS current type contributing the most. NR, non-responders. ***, P < 0.001; χ^2^ test. **(E)** Prevalence of RA, IA, SA and ultra-SA current components in siRNA-electroporated DRG neurons (top panels) and corresponding average maximal current amplitude (bottom panels). In the top panels, solid and striped colors indicate neurons with monophasic or biphasic MS currents, respectively. ***, P < 0.001; *, P < 0.05; χ^2^ test (top panels) and Kruskal-Wallis multiple comparison (bottom panels). **(F)** Stacked histogram showing the prevalence of RA, IA, SA and ultra-SA currents in IB4-positive neurons electroporated with control or *Tmem120a* siRNAs. Neurons responding with biphasic currents were classified according to the MS current type contributing the most. χ^2^ test shows no statistical difference. NR, non-responders. Inset: arrow shows a DRG neuron positive for IB4 (red signal) and siRNA transfection (green signal). Scale bar = 10 µm. **(G)** Prevalence of RA, IA, SA and ultra-SA current components in siRNA-electroporated IB4-positive neurons (top panels) and corresponding average maximal current amplitude (bottom panels). In the top panels, solid and striped colors indicate neurons with monophasic or biphasic MS currents, respectively. Mann-Whitney test shows no statistical difference. For all panels, n numbers are indicated in brackets. Error bars represent s.e.m.

We performed siRNA experiments in DRG neurons to determine whether *Tmem120a* could be responsible for a MS current. Since DRG neuron electroporation typically displays low transfection efficiency (≈10%), we assessed *Tmem120a* siRNA efficacy by RT-qPCR in NIH/3T3 electroporated cells (transfection efficiency 60-80%), which constitutively express *Tmem120a* (Fig.S4A). In these cells, *Tmem120a* siRNA transfection induced a decrease of 72 ± 5 % of *Tmem120a* mRNA level compared to non-targeting siRNA transfection (Fig.S4D). Interestingly, knock-down of *Piezo1* but not *Tmem120a* inhibited the MS current present in NIH/3T3 cells (Fig.S4F).

Co-electroporation of siRNAs and AcGFP1-plasmid was then performed in mouse DRG neurons, thereby allowing the fluorescent detection of transfected neurons. Non-targeting and *Piezo2* siRNAs were used as negative and positive controls, respectively. We first compared the proportions of DRG neurons classified according to their main MS current component. As previously reported (Coste et al., 2010), inhibition of *Piezo2* expression significantly (*p < 0*.*0001*, χ^2^ test) altered the relative proportion of DRG neurons exhibiting the different MS currents, with notable decrease in the incidence of RA current-expressing neurons from 38.1 to 10.0% and consequential increase of non-responsive neurons from 8.2 to 22.9%. By contrast, inhibition of *Tmem120a* expression had no effect on the relative proportion of MS currents compared to control conditions (*p = 0*.*9*, χ^2^ test) (Fig.5D). We next analyzed each MS current component quantified from mono- or bi-exponential fits of MS current recordings in siRNA-treated DRG neurons (Fig.5E). *Piezo2* siRNA treatment significantly reduced the proportion of neurons expressing the RA current component as well as the mean amplitude of the residual RA current. Once again, there were no significant changes in the proportion of any MS current components or in the mean amplitude of these components when *Tmem120a* expression was inhibited (Fig.5E).

To restrain the neuronal population investigated, we took advantage of the observation that TMEM120A protein immunostaining was detected in 80% of IB4^+^ DRG neurons (Beaulieu-Laroche et al., 2020). Therefore, we performed another set of experiments by selectively recording from IB4^+^ neurons identified using IB4-Alexa Fluor 568 conjugate staining. Consistent with our previous results, we found no differences regarding either the incidence or the amplitude of the different MS current components between *Tmem120a* and non-targeting siRNA-treated neurons (Fig. 5F-G). Furthermore, we cloned *Tmem120a* cDNA from mouse DRG into an expression vector. Cell-poking and pressure-clamp experiments showed that *Tmem120a* overexpression does not induce MS currents in HEK-P1KO (Dubin et al., 2017) transfected cells (Fig.S5). Altogether, these data suggest that *Tmem120a* does no contribute to any MS currents in DRG neurons.

### Inhibition of *Tmem150c* expression does not impair MS currents in DRG neurons

*Tmem150c* has been proposed to encode SA MS channels in proprioceptive DRG neurons contributing to motor coordination (Hong et al., 2016). This conclusion has been challenged by independent groups, because *Tmem150c* expression by itself does not appear to generate MS current but rather modulates the activity of mechano-gated channels including PIEZO1, PIEZO2 and the two-pore domain K^+^ channel TREK1 (Anderson et al., 2018; Dubin et al., 2017). Our scRNAseq data indicated that *Tmem150c* is not enriched in a particular subset of DRG neurons. Therefore, to gain more understanding about its role in MS current generation we recorded from *Tmem150c* siRNA electroporated DRG neurons. We found no differences whatsoever regarding either the incidence or the amplitude of the different MS current components in *Tmem150c* siRNA-treated neurons compared with non-targeting siRNA-treated neurons (Fig. 5D-E). These data argue against a role of *Tmem150c* in sustaining a particular type of MS current in DRG neurons.

### Genes associated with DRG MS current types

Enrichment of *Piezo2* in the subset of RA current expressing neurons provides a rationale to identify candidate genes coding for or regulating MS channels. Therefore, we looked for transcripts differentially expressed (P < 0.05) and enriched at least 2-fold in neurons grouped by MS current type (Fig.6A-D). We provide a list of transcripts associated with RA, IA, SA and ultra-SA currents containing 212, 404, 125 and 84 protein coding genes, respectively (Table S2). As can be seen in figure 6E, *Piezo2* emerged in the top 15% of most expressed genes amongst the enriched ones. Therefore, we reasoned that both enrichment and mRNA abundance in the corresponding neuronal population may be good criteria to pinpoint molecular determinants associated with MS current phenotype. Additionally, gene ontology (GO) analysis was used to annotate these genes with relevant knowledge such as detection of a mechanical stimulus, ion channel activity and integral component of the membrane. The top 40 of the most expressed genes associated with MS current type are listed in figure 6E. Among those, *Piezo2* is the only gene known to encode a protein with demonstrated function in mechanical transduction. Moreover, the few genes known to encode ion channels, such as *Scn10a* encoding the voltage activated sodium channel Nav1.8 (Han et al., 2016), are well characterized and unlikely to encode MS channels. Several genes are known to code for proteins inserted in the membrane, a feature common to ion channels. However, as GO resource reflects our current biological knowledge many of these candidate genes remain to be functionally characterized. By refining from 13,000 transcripts expressed per DRG neuron to only hundreds of candidates, this dataset represents a valuable resource for the identification of genes involved in the generation of DRG MS currents.

**Figure 6.**
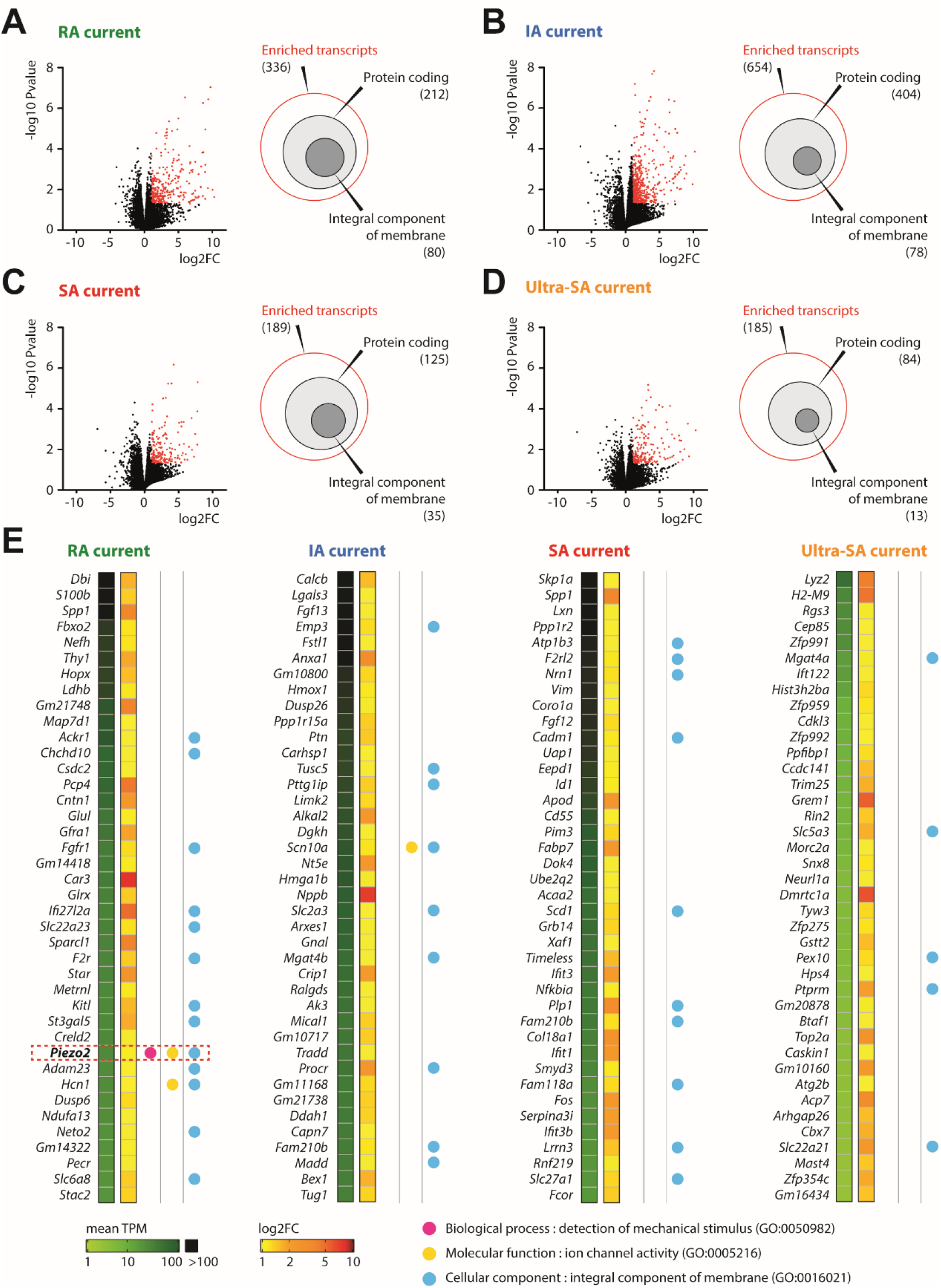
Potential molecular determinants of DRG MS channels. (**A-D**) Left panels: Volcano plots of transcripts expressed in neurons grouped by MS current type, as indicated. Red dots illustrate transcripts enriched (p < 0.05) with FC > 2, the full list of which is available in Table S2. Right panels: Euler diagrams showing among the enriched transcripts those that code for proteins (light gray) and those that code for integral membrane proteins (dark gray) (**E**) Lists of the top 40 most expressed protein-coding genes among enriched transcripts in DRG neurons grouped by MS current type. GO analysis was used to annotate genes.

## DISCUSSION

### Patch-clamping of cultured DRG neurons is compatible with scRNAseq classification

Direct relationship between mechanosensory properties and molecular phenotypes of DRG neurons determined by whole transcriptome analysis has never been explored. To identify molecular features associated with specific mechanosensory functions and phenotypes, we have combined scRNA-seq profiling with electrophysiological recordings of individual DRG neurons. We provided full transcriptome data from single neurons in culture after characterization of MS currents, which were classified based on their inactivation kinetics. Coupling patch-clamp recording with mechanical stimulation required the use of adhering cultured DRG neurons. This approach is labor-intensive and low throughput and differs somehow from previous scRNAseq classification studies for which DRG neurons were collected shortly after ganglionic dissociation (Li et al., 2016; Sharma et al., 2020; Usoskin et al., 2015; Zeisel et al., 2018). Yet, 85% of the processed mRNA samples met quality standards for RNAseq. The molecular classification of our small and heterogeneous group of DRG neurons however was challenging because of the resulting low statistical power. Advances in microfluidics for single cell partitioning have enabled scRNAseq studies performed on samples of over one thousand cells, subdividing DRG neurons into a total of up to 17 genetic clusters (Sharma et al., 2020; Zeisel et al., 2018). To achieve high-quality classification of our neuron sample, we combined our single-cell transcriptomes with the larger single-DRG neuron dataset from Zeisel and collaborators (Zeisel et al., 2018). Therefore, despite its modest size, we succeeded to map all the neurons from our sample into DRG neuron genetic clusters. They are distributed among 11 of the 17 genetic clusters corresponding to 7 of the 8 main classes of neurons. The clusters which are not represented are PSNF2 (from NF neurons), PSNP2 (from NP1 neurons), PSPEP4 (from PEP1 neurons) and the three clusters PSPEP6-8, which together constitute the TrpM8 population. The absence of TrpM8 neurons in our sample is consistent with their lack of role in mechanosensation (Knowlton et al., 2013). It is possible that PSNP2 and PSPEP4 constitute subgroups of NP1 and PEP1 neurons that are not MS, but our relatively small sample of neurons does not allow a definite conclusion. However, the absence of PSNF2 neurons, thought to represent a cluster of proprioceptors (Usoskin et al., 2015) likely stems from an undersized sample. Yet, our results attest that the culture of neurons, coupled to patch-clamp recording, does not prevent their genetic classification and offers an extensive view of the main populations of DRG MS neurons. Thus, patch-clamp recording of MS currents is compatible with the production of a quantitative dataset that resolves mRNAs for all known genes in single mechanosensory neurons.

### Distribution of MS current types in genetically defined neurons

The distribution of MS currents in DRG neurons has been addressed over the past by distinguishing neuronal subsets using morphological and electrophysiological parameters as well as IB4 staining (Coste et al., 2007; Drew et al., 2004; Drew et al., 2002; Hu and Lewin, 2006; Lechner and Lewin, 2009; Prato et al., 2017). These approaches led to the view that large diameter, non-nociceptive neurons express mainly RA MS currents whereas medium- and small-diameter nociceptors express MS currents with slower kinetics. Subsets of DRG neurons labeled *in vivo* using genetically modified mice confirmed the tendency of proprioceptors and non-nociceptive A-type LTMRs to express RA MS currents (Woo et al., 2015; Zheng et al., 2019), and showed that C-LTMR express MS currents with rapid and/or slow kinetics (Delfini et al., 2013; Lou et al., 2013). The accurate characterization of four distinct types of DRG MS currents in classified neurons using scRNAseq enabled to study their distribution in DRG neuron subsets.

Our results showed that the RA MS current is the main subtype in NF neuronal population, in line with the remarkable phenotype of *Piezo2* KO animals related to innocuous touch and proprioception (Florez-Paz et al., 2016; Ranade et al., 2014b; Woo et al., 2015).

Consistently with *Piezo2* modest contribution to mechanical nociception (Murthy et al., 2018b), RA current type contributes to a lesser extent to MS currents in NP1, PEP1 and PEP2 neuronal subpopulations. The PEP1 neurons express predominantly the IA MS current type. This suggests that IA MS current may be the main transducer of noxious mechanical stimuli in C-type peptidergic nociceptors. The contribution of SA and ultra-SA currents in these neurons seems minimal. Interestingly, the opposite is true in PEP2 neurons, in which ultra-SA current type contributes the most, whereas IA current type is almost absent. This suggests that Aδ-nociceptors and C-type peptidergic nociceptors use distinct sets of MS ion channels for mechanical pain detection. These differences could contribute to the signaling of distinct mechanosensory modalities by PEP1 and PEP2 nociceptors.

Importantly, RA current type is the main contributor of MS currents in NP2 neurons. These neurons, which have been shown to respond to mechanical stimulus, heat, and a large array of pruritogens, mediate itch behavior but not pain (Han et al., 2013). Therefore, these results suggest that *Piezo2* actively contributes to itch sensing, by opposition to its expression in Merkel cells in which *Piezo2* inhibition produces alloknesis, a pathological itch sensation produced by innocuous touch (Feng et al., 2018).

### No evidence for *Tmem120a* and *Tmem150c* in sustaining slow MS currents

While a compelling body of evidence indicates that *Piezo2* generates RA-type MS current in DRG neurons, the molecular identity of genes coding for slow MS currents has remained elusive (Delmas and Coste, 2013; Ranade et al., 2015). Recently, new gene candidates have been proposed to encode slow-type MS currents in DRG neurons. *Tmem120a* has been suggested to be expressed in small-to medium-diameter neurons, and to mediate the ultra-slowly adapting MS current (Beaulieu-Laroche et al., 2020). Our results do not support this conclusion. First, our single-cell transcriptome analysis indicates that *Tmem120a* expression is not restricted to nociceptors and is not enriched in slowly or ultra-slowly adapting MS current-expressing neurons. This is consistent with our analysis of the cross-sectional area distribution of *Tmem120a*-positive DRG neurons in *ISH* experiments. When normalized to the overall population, *Tmem120a* was detected in all subtypes of DRG neuron cell body diameters, including the largest ones expressing NF200. Nevertheless, the possibility that *Tmem120* contributes to MS currents without being selectively expressed/enriched in a particular population was still open. However, our *Tmem120a* siRNA experiments performed either on the whole DRG neuron population or restricted to IB4^+^ neurons showed no changes in the proportion of slowly or ultra-slowly MS current-expressing neurons, nor in the mean amplitude of these current components. Our siRNA assays beneficiated from our extensive kinetics classification of MS currents and from the inclusion of a *piezo2* siRNA effective target knockdown. Thus, the broad expression of *Tmem120a* in DRG neurons, along with the lack of appreciable effects of *Tmem120a* siRNA treatment on endogenous MS currents, cast doubt about its role in mediating SA/ultra-SA MS currents. Moreover, overexpression of mouse *Tmem120a* in heterologous system did not induce MS channel activity. However, our data do not question *Tmem120a* involvement in pain sensing, which has been shown recently in inflammatory, but not neuropathic, mechanical hyperalgesia in rats (Bonet et al., 2020). The molecular function of *Tmem120a* needs to be clarified in future work.

*Tmem150c* has been proposed to encode slowly adapting MS channels in DRG proprioceptive neurons (Hong et al., 2016) but this inference has been challenged. Indeed, we and others have shown that, contrary to what was initially reported, *Tmem150c* expression in heterologous system does not induce MS currents by itself (Anderson et al., 2018; Dubin et al., 2017), but instead modulates the properties of MS ion channels when co-expressed (Anderson et al., 2018). In addition, no slowly adapting current type was detected in proprioceptive neurons labeled *in vivo* using genetically modified mice (Woo et al., 2015). *Tmem150c* was found to be broadly expressed in our single-cell transcriptome analysis, and our siRNA experiments showed that *Tmem150c* knock-down caused no changes in the proportion or amplitude of any MS currents, suggesting that *Tmem150c* is not a prime contributor to MS currents in DRG neurons. To note, *Tmem150c* has been proposed recently to contribute to slowly adapting MS currents in nodose ganglion baroreceptors (Lu et al., 2020). Future studies will help resolve these controversies.

### Molecular bases of the kinetically distinct MS current subtypes in DRG

Intensive efforts have been deployed to identify mammalian MS channels for decades, since the earliest indications of their existence in auditory hair cells (Corey and Hudspeth, 1979). Identifying MS channel genes represents a crucial challenge for the field of nociception and may have many clinical implications in acute as well as inflammatory and chronic pain. In addition, it would be potentially relevant beyond somatosensation as exemplified by PIEZO channel implication in many physiological functions and diseases in humans (Ma et al., 2018; Ma et al., 2021; Marshall et al., 2020; McMillin et al., 2014). Therefore, by correlating DRG MS current properties with co-expressed genes at the single cell level, our study provides a valuable resource for identifying new molecular entities involved in mechanotransduction.

Our extensive characterization of MS current properties from large neuronal samples showed that 4 distinct types of MS currents are present in mouse DRG neurons. It is now well established that *Piezo2* encodes MS channels responsible for RA currents. Could *Piezo2* contribute to other types of MS currents? It has been shown that overexpression of *Tmem150c* with *Piezo2* in heterologous systems induces slowing of PIEZO2 current inactivation to values close to IA current type (Anderson et al., 2018). However, *Piezo2* deletion or knock-down studies in DRG neurons, including this work, have shown no alteration of IA currents (Coste et al., 2010; Ranade et al., 2014b), and a possible compensatory increase in IA current responses has been reported in proprioceptors (Woo et al., 2015). This suggests that *Piezo2* does not encode MS channels responsible for IA currents in DRG neurons. The situation differs in trigeminal corneal neurons where a non-specific decrease of currents with intermediate and slow kinetics has been reported in *Piezo2* KO mice (Fernandez-Trillo et al., 2020). We also observed a modest effect of *Piezo2* inhibition on the amplitude of ultra-SA currents. It cannot be excluded that this decrease is due to a RA current component masked in neurons with large ultra-SA currents. However, this modest impairment is still observable when current amplitude was measured at the end of the mechanical stimulus, *i*.*e*. when RA currents are expected to be fully inactivated (data not shown). Inhibition of a *Piezo2* splice variant could be responsible for this effect, as *Piezo2* extensive alternative splicing affecting inactivation kinetics has been reported in DRG neurons, although Piezo2 splicing variants displayed inactivation kinetics still in the range of RA current subtype (Szczot et al., 2017). Whatsoever, this effect is slight and does not alter the proportion of neurons expressing the ultra-SA current, arguing that the primary contributor of ultra-SA currents is distinct from PIEZO2.

Notwithstanding, the nature of the molecular entities involved in IA-, SA- and ultra SA-currents remains unknown. In addition, the question of whether these currents are generated by multiple or single molecular entities is not yet resolved. Indeed, proximate environment such as membrane composition, attachment to extracellular matrix or subcortical cytoskeleton organization can potentially modify membrane bilayer tension and inserted MS channels properties including kinetics (Cox et al., 2016; Syeda et al., 2016; Teng et al., 2015). In this scenario, accessory subunits conferring distinct inactivation properties to a unique MS channel could be encoded by genes found to be enriched in a subset of DRG neurons.

Evidence suggests that the different currents are sustained by molecularly distinct ion channels. Indeed, three types of cationic MS ion channels with sustained activity but distinguishable from their unitary conductance have been described in DRG neurons (Cho et al., 2006; Cho et al., 2002). Moreover, ultra-SA currents are blocked selectively by the conopeptide NMB1 (Drew et al., 2007), suggesting that this current is sustained by a specific pool of MS channels. Hence, several MS channel coding genes remain to be discovered in DRG neurons. By identifying the genes enriched in DRG neurons with distinct mechanical phenotypes, our study lays the foundations for the identification of their molecular entities. Testing these candidates using siRNA experiments coupled with patch-clamp recording in DRG neurons should identify genes involved in mechanosensation. Importantly, given the broad involvement of mechanotransduction in physiological functions and the dynamism of the mechanobiology field, illustrated by the development of novel mechanobiology assays (Kurth et al., 2012; Xu et al., 2018), our data could also be used to identify mechanosensors or associated proteins in other biological tissues.

## Acknowledgements

This project has received funding from the European Research Council (ERC) under the European Union’s Horizon 2020 research and innovation programme (Grant agreement No. 678610). RNA sequencing was done by Profilexpert, “Génomique & Microgénomique” service, Lyon 1 University, SFR santé Lyon-Est, UCBL-Inserm US 7-CNRS 3458, 8 av. Rockefeller, 69373 Lyon Cedex 8, FRANCE. We thank Virginie Penalba and Axel Fernandez for expert technical assistance, and Dr. Nancy Osorio, Dr. Françoise Padilla and Dr. Nicolas Grillet for critical reading of the manuscript.

## Author contributions

L.B. and B.C. recorded and collected DRG neurons for RNA sequencing. N.S. and Y.M. prepared sequencing libraries. T.P. conducted all bioinformatics analysis. N.S. performed RNAscope experiments. A.L. conducted qPCR experiments. T.P. and L.B. conducted siRNA experiments. B.C. conceived and supervised all aspects of the project. T.P. and B.C. prepared the figures and T.P., P.D. and B.C. wrote the manuscript with input from L.B.

## Declaration of interests

The authors declare no competing interests.

## STAR METHODS

### KEY RESOURCES TABLE

| REAGENT or RESOURCE | SOURCE | IDENTIFIER |
| --- | --- | --- |
| **Antibodies** |  |  |
| Anti-Peripherin Antibody | Sigma-Aldrich | #AB1530 |
| Anti-NFH (NeuroFilament Heavy chain) antibody | AVES Labs | #NFH |
| Alexa Fluor® 647 AffiniPure Donkey Anti-Rabbit IgG (H+L) | Jackson ImmunoResearch | #711-605-152 |
| Rhodamine (TRITC) AffiniPure Donkey Anti-Chicken IgY (IgG) (H+L) | Jackson ImmunoResearch | #703-025-155 |
| **Bacterial strains** |  |  |
| pIRES2-AcGFP1 | Clontech | #632435 |
| **Critical Commercial assays** |  |  |
| RNAscope fluorescent multiplex reagent kit | Advanced Cell Diagnostics | #323133 |
| RNAscope Protease IV Reagent | Advanced Cell Diagnostics | #322340 |
| Fluorescent Multiplex Detection Reagents | Advanced Cell Diagnostics | #320851 |
| Wash Buffer Reagents | Advanced Cell Diagnostics | #310091 |
| RNAscope 3-plex Negative Control Probe | Advanced Cell Diagnostics | #320871 |
| *Tmem120a* probe | Advanced Cell Diagnostics | #513211 |
| SMART-seq v4 Low Input Kit | Clontech | #634888 |
| NucleoSpin RNA Mini kit | Macherey-Nagel | #740955 |
| DNA-free DNase Treatment | Ambion | #AM1906 |
| iScript™ Reverse Transcription Supermix for RT-qPCR | Bio-Rad | #1708840 |
| KAPA SYBR® FAST qPCR Kit Master Mix (2X) Universal | Kapa Biosystems | #KK4600 |
| Neon™ Transfection System 10 µL Kit | Invitrogen | #MPK1096 |
| **Others** |  |  |
| Optimum Cutting Temperature medium | CellPath Ltd | #KMA-0100-00A |
| Phosphate buffered saline (PBS) | Sigma-Aldrich | #P4417 |
| isoflurane | IsoVet | #200265 |
| sucrose | Sigma-Aldrich | #S0389 |
| 16% Formaldehyde (w/v), Methanol-free | ThermoFisher Scientific | #PI28908 |
| PIPES | Sigma-Aldrich | #P6757 |
| HEPES | Sigma-Aldrich | #H3375 |
| EGTA | Sigma-Aldrich | #E4378 |
| Magnesium chloride hexahydrate (MgCl<sub>2</sub>) | Sigma-Aldrich | #M2670 |
| Triton™ X-100 | Sigma-Aldrich | #T8787 |
| Gelatin from cold water fish skin | Sigma-Aldrich | #G7765 |
| ProLong™ Gold Antifade Mountant | ThermoFisher | #P36930 |
| Thermo Scientific™ SuperFrost Plus™ Adhesion slides | Fisher Scientific | #J1800AMNZ |
| Diethyl pyrocarbonate (DEPC) | Sigma-Aldrich | #159220 |
| Dulbecco's Modified Eagle's Medium (DMEM) High glucose, pyruvate | ThermoFisher | #41966-029 |
| Fetal Bovine Serum (FBS) | Gibco | #10270-098 |
| Pen Strep (Penicillin Streptomycin) | Gibco | #15140-122 |
| Dulbecco's Phosphate Buffered Saline (DPBS) | Gibco | #14190-094 |
| GelRed Nucleic Acid Gel Stain, 10,000X | Biotium | #41003 |
| ExactGene 100bp ladder | Fisher BioReagents™ | BP2571100 |
| **Devices** |  |  |
| Neon® Transfection System | Invitrogen | #MPK5000 |
| Applied Biosystems 7500 Fast Real-Time PCR System | Applied Biosystems |  |

### CONTACT FOR REAGENT AND RESSOURCE SHARING

Further information and requests for reagents and resources should be directed to and will be fulfilled by the Lead Contact, Bertrand Coste.

### EXPERIMENTAL MODEL

#### Mice

Male mice (C57Bl/6J background) were 10-12 weeks old. All animals were used in accordance with the European Community guidelines in the care and use of animals (2010/63/UE). This project was approved by the Institutional Review Board of the regional ethic committee.

### METHOD DETAILS

#### Dorsal root ganglion neuron culture

Mice were anesthetized with isoflurane and killed by decapitation. DRG were isolated and incubated in enzyme solution containing 2 mg.ml^-1^ of collagenase 1A (Sigma-Aldrich) and 5 mg.ml^-1^ of dispase II (ThermoFisher) for 45 min at 37 °C. The tissue was washed several times and triturated in Hanks’ balanced salt solution (HBSS, Gibco). The resulting suspension was centrifuged (300×g for 5 min) in 1 ml HBSS and 1 ml Bovine Serum Albumin (BSA, 15%, Sigma-Aldrich). The pellet was rinsed with Dulbecco’s Phosphate Buffered Saline (DPBS, Gibco) and centrifuged (300×g for 2 min). Cells were then plated onto poly-D-lysine-coated 12-mm round glass coverslips (Corning) in 24-well plates and were grown in Dulbecco’s modified Eagle’s medium (DMEM, High glucose, pyruvate, ThermoFisher) supplemented with 10% heat-inactivated Fetal Bovine Serum (FBS, Gibco), 100 U.ml^-1^ (1%) penicillin– streptomycin (Gibco), 100 ng.ml^-1^ nerve growth factor (NGF, Merk Millipore), and 50 ng.ml^-1^ glial-derived neurotrophic factor (GDNF, ThermoFisher). Cells were recorded at 3-5 days *in vitro* (DIV).

#### Electrophysiology

Patch-clamp experiments were performed under whole-cell configuration using an Axopatch 200B amplifier (Axon Instruments). Patch pipettes had resistances of 2-3 MΩ when filled with an internal solution consisting of (in mM) 140 CsCl, 10 Hepes, 5 EGTA, 1 CaCl_2_, 1 MgCl_2_, 4 MgATP and 0.4 Na_2_GTP (pH adjusted to 7.3 with CsOH). Internal solution was supplemented with 0.4 U/µl of RNaseOUT (Invitrogen) for cell harvesting experiments. The extracellular solution consisted of (in mM) 133 NaCl, 3 KCl, 1 MgCl_2_, 10 Hepes, 2.5 CaCl_2_, 10 glucose (pH adjusted to 7.3 with NaOH). All experiments were done at room temperature. Currents were sampled at 20 kHz and filtered at 2 kHz.

#### Mechanical stimulation

Mechanical stimulation was achieved using a firepolished glass pipette (tip diameter 3-4 μm) positioned at an angle of 80° and in contact with the cell being recorded. Downward movement of the probe toward the cell was driven by a Clampex controlled piezo-electric crystal microstage (E625 LVPZT Controller/Amplifier, Physik Instrumente). The probe had a velocity of 0.7 μm.ms^-1^ during the ramp segment of the command for forward motion, and the stimulus was applied for 150 ms. Inward MS currents were recorded at a holding potential of -80 mV.

MS currents were characterized by applying series of 0.5 μm incremental steps every 10 s. This was done up to patch rupture, except for neurons dedicated to scRNAseq experiments. For this set of experiments, mechanical stimulation of neurons was stopped after MS currents with a peak amplitude of more than 100 pA were elicited in at least two consecutive incremental steps, preserving the integrity of the cellular content before the harvesting. The 3 non-responsive neurons included in our scRNAseq sample have been mechanically stimulated up to 10 µm probe displacement without disrupting the patch.

#### Mechanosensitive current analysis

Clampfit 10.7 (Molecular Devices) software was used to analyze recordings and biophysical parameters. The decay of inactivation from the peak of current was fitted with mono or bi-exponential function of the form 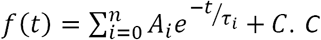 was set to the baseline value of current once the mechanical stimulus is removed. Inactivation time constant *τ* was used to identify the type of MS current components. To note, *τ* measurements of ultra-SA currents are approximave due to the relative short lasting (150 ms) of the mechanical stimulus. When fitted with bi-exponential equation, the contribution (in percentage) of each component to the whole current is determined from the ratio of *A*_*i*_ to the amplitude of the peak current (see Fig.S2B). Recordings of MS currents displaying no clear deactivation once the mechanical stimulus is removed were discarded. We arbitrarily set a component detection threshold at 10% of the peak current, *i*.*e*. so that only detected components contributing to at least 10% of MS currents were included into analyzes, to avoid mis-identification of current components.

#### Pressure-clamp experiments

Pressure-clamp experiments were performed under cell-attached configuration. Patch pipettes had resistances of 2-3 MΩ when filled with a solution consisting of (in mM) 130 NaCl, 5 KCl, 10 HEPES, 1 CaCl_2_, 1 MgCl_2_, 10 TEA-Cl (pH 7.3 with NaOH). External solution used to zero the membrane potential consisted of (in mM) 145 KCl, 10 HEPES, 1 MgCl_2_, 10 glucose (pH 7.3 with KOH). Currents were sampled at 20 KHz and filtered at 2 KHz. Membrane patches were stimulated with negative pressure pulses through the recording electrode using a Clampex controlled pressure clamp HSPC-1 device (ALA-scientific). Stretch-activated channels were recorded at a holding potential of -80 mV.

#### Single-cell library preparation

After electrophysiological recording, the cytoplasmic content of the neuron was harvested into the patch pipette, transferred into lysis solution (RNaseOUT 8 U.µl-1, 2% Triton X-100) and flash frozen. Reverse transcription was performed using a SMART-seq v4 Low Input Kit (Clontech) directly on the cell lysate according to the manufacturer’s protocol. After cDNAds amplification, 50 μl of the sample were subjected to cDNA purification on AMPure XP beads. 0.5 ng of each purified cDNAds were used to construct the sequencing library using Nextera XT kit (Illumina) with fragments over 300-bp length from each neuron.

#### Processing, quality control and filtering of single-cell RNA-seq data

cDNAds libraries were sequenced either in a NextSeq 500 (Illumina) using paired-end reads (76 pb) with an average depth of 29 million reads per cell. The raw reads (FastQ files) were cleaned by removing adaptor sequences, short sequences (length < 25 bp), low-quality bases (quality score < 30) and ambiguous sequences (i.e., reads with more than 30% unknown bases ‘N’) using CutAdapt 1.9.1 software (Martin, 2011). TopHat-2 v2.1.0 was used to map the cleaned RNA-seq reads to the mouse genome (mm10 / GRCm38.90) with two mismatches, two gaps and one multihit allowed. TopHat-2 splicing algorithm was used to map reads covering splice junctions, thereby improving the utilization of reads. After genome mapping, gene counts matrix was determined using GenomicAlignment (version 1.24.0) (Lawrence et al., 2013) using GTF annotation. A set of mitochondrial genes (mt-Atp6, mt-Atp8, mt-Co1, mt-Co2, mt-Co3, mt-Cytb, mt-Nd1, mt-Nd2, mt-Nd3, mt-Nd4, mt-Nd4l, mt-Nd5, mt-Nd6) were used to check viability of each sample: the sample is considered as non-apoptotic if the expression of these genes is less than 10% of total count in the same cell. Read counts were analyzed in the R/Bioconductor environment (version 3.12; www.bioconductor.org) with the R package edgeR. The gene expression value was normalized between samples by TMM with a gene detection threshold of 20 counts in at least two cells. For the generation of heat maps shown in Figures 2B, z-transformed TMM count data was used. Relative expression levels were obtained by sample normalization using TPM (transcripts per kilobase million).

#### Dorsal root ganglion neuron clustering

The mapping of the scRNAseq neurons in DRG neuron genetic clusters was done by a cross-dataset normalization approach. We used Mutual Nearest Neighbours (MNN) correction (Haghverdi et al., 2018) implemented in the R package batchelor to combine our dataset with Zeisel and collaborators scRNAseq sensory neuron dataset (Zeisel et al., 2018). Briefly, a principal component analysis (PCA) was performed on the 211 most highly variable genes in DRG neurons (most highly enriched genes for each cluster) (Zeisel et al., 2018) to correct for batch effect. Identification of MNN was done in this reduced dimension space using the following parameters: dimension = 60 and k = 10. To assign each individual neuron to a specific cluster, we built a k nearest-neighbors (kNN) graph with k = 28 using Euclidean distance. Individual neurons were first assigned to one of the main cluster: NF, NP or PEP. Next, we repeated the procedure (PCA implemented in MNN correction, kNN clustering) for each main cluster. We were able to assign each cell to a specific sub-cluster: PSNF1, PSNF2, PSNF3, PSNP1, PSNP2, PSNP3, PSNP4, PSNP5, PSNP6, PSPEP1, PSPEP2, PSPEP3, PSPEP4, PSPEP5, PSPEP6, PSPEP7 or PSPEP8. Corrected expression values obtained during this procedure were only used for cluster assignment and visualization with t-distributed stochastic neighbor embedding (tSNE) projection (perplexity = 30, max iteration = 1000). Downstream analysis was performed using custom R scripts.

#### Electroporation of DRG neurons

DRG cultures were electroporated using the Neon® Transfection System (Invitrogen) according to the manufacturer’s protocol. After the centrifugation prior to cell plating, the cell pellet was resuspended in Neon® Transfection System resuspension Buffer R (Invitrogen) and gently mixed with 30 ng.µl^-1^ of pIRES2-AcGFP1 plasmid (Clontech) and 250 nM siRNA pool. Ten microliters of the cell suspension were then electroporated with the following program: 1200 V, 2 pulses, 20 ms. Electroporated cells were then plated onto poly-D-Lysine-coated 12-mm round glass coverslips in 24-well plates in antibiotics-free media. Three hours later, the medium was replaced by DMEM supplemented with 10% heat-inactivated FBS, 1% penicillin–streptomycin, 100 ng.ml^-1^ NGF and 50 ng.ml^-1^ GDNF. Pools of 4 siRNA targeting *Piezo2* or *Tmem120a*, or non-targeting siRNA for control experiments were purchased at Horizon Discovery (#L-163012-00-0005, #L-040281-01-0005 and #D-001810-10-05 for *Piezo2, Tmem120a* and control siRNA, respectively) and the pool of 4 siRNA targeting *Tmem150c* was purchased at Qiagen (#GS231503). Media were changed 48 hours later. Transfected DRG neurons were visualized by the expression of GFP. For experiments performed on IB4-positive neurons, neurons were incubated with 1 µg.ml^-1^ isolectin GS-IB4 from Griffonia simplicifolia, AlexaFluor 568 conjugate (Invitrogen) at 37°C for 10 minutes before recording.

#### NIH/3T3 cell culture and electroporation

NIH/3T3 cells were grown in DMEM supplemented with 10% heat-inactivated FBS (Gibco) and 100 U.ml^-1^ (1%) penicillin–streptomycin (Gibco). Cell passage was done when cells reached 80% confluency. For electroporation experiments, the Neon® Transfection System (Invitrogen) was used according to the manufacturer’s protocol. Briefly, prior to cell plating, the cell pellet was resuspended in Neon® Transfection System Buffer R at a cell density of 1 ×10^7^ cells.ml^-1^. 250 nM siRNA was added to 10 µl of cell suspension and electroporation was done with the following program: 1400 V, 1 pulse, 30 ms. Electroporated cells were then transferred directly into 24-well tissue culture plates (1×10^5^ cells per well) for RNA extraction, or onto poly-D-lysine-coated 12-mm round glass coverslips for patch-clamp recording experiments 72 hours later.

#### Molecular cloning of *Tmem120a*

Primers were designed from cDNA sequence of *Tmem120a* from the NCBI database (NM_172541). An 1.096 kb fragment was amplified from cDNA libraries generated from adult C57BL/6J DRG total RNA using primers fwd (5’ GATTATGCATGCCGTGGACAAAGACATGCAGT 3’) and rev (5’ GATTATGGATCCTCAGTCCTTCTTGTTCCCGT 3’) and cloned into pIRES2-AcGFP1 vector (Clontech, #632435) with NheI and BamHI restriction sites. The protein sequence of *Tmem120a* that we cloned is identical to NCBI Reference Sequence NP_766129.

#### HEK-P1KO cell culture and transfection

PIEZO1-deficient HEK293T cells (Dubin et al., 2017) were grown in Dulbecco’s Modified Eagle Medium containing 4.5 mg.ml^-1^ glucose, 10% (vol/vol) FBS, 100 U.ml^-1^ (1%) penicillin/streptomycin. Cells were plated onto poly-D-lysine-coated 12-mm round glass coverslips (Corning) in 24-well plates and transfected using lipofectamine 2000 (Invitrogen) according to the manufacturer’s instruction. Each well was transfected with 600 ng.ml^-1^ of specified constructs. GFP positive cells were recorded 48 hours later.

#### qPCR experiments

Electroporated NIH/3T3 cells were harvested after 48 hours of incubation at 37°C in antibiotics-free growth medium. Total RNA was extracted using NucleoSpin RNA Mini kit (Macherey-Nagel) according to the manufacturer’s instructions. DNA-free DNase Treatment (Ambion) was additionally used to remove genomic DNA contamination and iScript™ Reverse Transcription Supermix for RT-qPCR (Bio-Rad) was employed for cDNA synthesis. RT-cDNA samples generated with the iScript No-RT Control Supermix included in the kit were also synthesized for each sample as an additional verification step for the absence of genomic DNA contamination.

Quantitative PCR (qPCR) was performed on an Applied Biosystems 7500 Fast Real-Time PCR System (Applied Biosystems) using KAPA SYBR^®^ FAST qPCR Kit Master Mix (2X) Universal (Kapa Biosystems). A 12 µl reaction mix was made using 4 µl of diluted cDNA template (1 ng/µl), specific primers designed for *Tmem120a* as well as reference genes (200 nM each), ROX Reference Dye Low (0.24 µl) and SYBR Green I Master Mix (6 µL) as recommended by the manufacturer. Cycling conditions were as follows: 95°C for 3 seconds, then 60°C for 30 seconds, 40 cycles. Reactions were performed in duplicate and melting-curve analysis was performed to assess the specificity of each amplification.

For each sample, Ct values obtained for gene of interest (GOI) were normalized in relation to the average Ct values of all reference genes (RG) (ΔCt = Ct_GOI_ – Mean Ct_RG_). The difference between the ΔCt of the sample of interest and the control sample (ΔΔCt = ΔCt_sample of interest_ – ΔCt_control sample_) was then calculated, and finally the 2−ΔΔCt value to express the results as relative expression levels of gene of interest in samples of interest versus the control sample (non-targeting siRNA).

#### Gel electrophoresis of RT-qPCR products

RT-qPCR amplicons produced in the presence or absence of RT enzyme were size-separated by electrophoresis on a 3.5% agarose gel stained with GelRed (Biotum), in comparison with ExactGene 100 bp ladder (Fisher BioReagents™) and photographed under UV light. The expected amplicon sizes are 144 bp for *Tmem120a*, 100 bp for *Piezo1*, 99 bp for *Piezo2*.

#### Primers

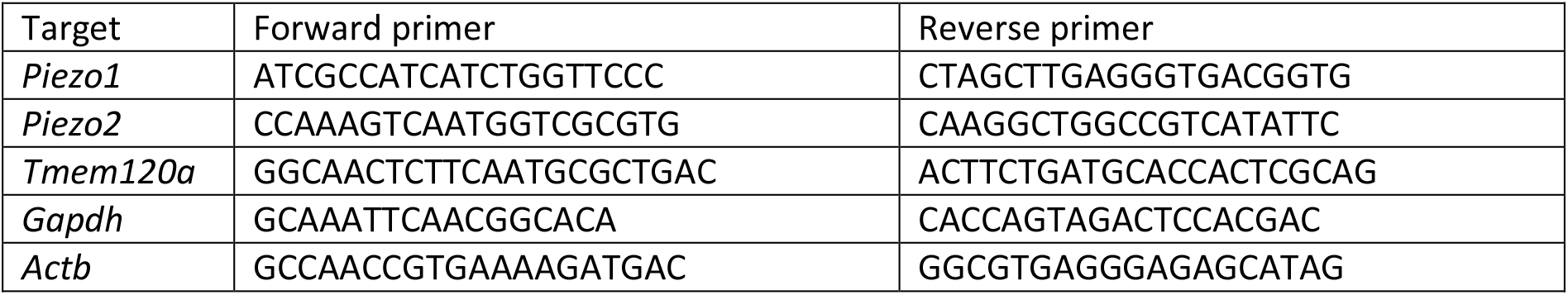

#### *In Situ* Hybridization experiments

Mice were anesthetized with isoflurane and killed by decapitation. Thoraco-lumbar vertebral spines were freshly collected, cut transversely, and transferred to a 4% sucrose solution (in 1X PBS) during 1h30 at 4°C, then to a 20% sucrose solution (in 1X PBS) overnight at 4°C. Then, DRG were isolated and embedded in Optimum Cutting Temperature medium (OCT; CellPath Ltd; #KMA-0100-00A) and cut into 14 µm sections with a CM350 S cryostat (Leica). *In situ* hybridization (ISH) was carried out using the RNAscope fluorescent multiplex reagent kit (Advanced Cell Diagnostics, a Bio-Techne brand, United States; #323133), according to the manufacturer’s instructions. Briefly, tissue sections were post-fixed with 4% paraformaldehyde during 30 min at 4°C and rinsed with 1X PBS. After ethanol dehydration and a Protease IV pretreatment for 30 minutes at room temperature, sections were incubated with a mouse *Tmem120a* probe (ACD; #513211, accession number #NM_172541.2) for 2h at 40°C. The negative control, designed by ACD Bio, contains probes targeting the *tapB* gene from the *Bacillus subtilis* strain SMY (ACD; #320871, accession number #EF_191515). The signal was detected and amplified with a succession of four pre-amplifiers supplied in the kit.

#### Co-immunohistochemistry experiments

NF-200 and peripherin immunostainings were done following ISH. After washing for 10 min at room temperature with PHEM buffer (consisting of in mM: 60 Pipes, 25 Hepes, 10 EGTA, 2 MgCl_2_) supplemented with 0.1% Triton X-100, non-specific sites were blocked with a reagent containing 3% Gelatin from cold water fish skin (Sigma-Aldrich; #G7765). Samples were incubated with primary antibodies against peripherin (1:200; rabbit polyclonal; Sigma-Aldrich; #AB1530) and NF-200 (1:1000; chicken polyclonal; AVES Labs; #NFH) overnight at 4°C. Primary antibodies were detected by using secondary antibodies conjugated to Alexa Fluor 647 (1 :1000 ; Donkey anti-rabbit polyclonal IgG ; Jackson ImmunoResearch, #711-605-152) and Alexa Fluor 568 (1:1000; Goat anti-chicken polyclonal IgY; Invitrogen, #A-11041), respectively, incubated for 1 h at room temperature. Sections were rinsed with PHEM buffer, dried and mounted in Prolong Gold Antifade Mountant (ThermoFisher; #P36930). Images were acquired using a LSM 780 laser-scanning confocal microscope (Zeiss) with Ar and He-Ne lasers using a 63X oil-immersion objective and treated with ImageJ software.

#### Statistical analysis

Unless mentioned otherwise in Method Details above, data were analyzed using GraphPad Prism 8.2.0 (San Diego, USA). Statistical details can be found in the figure legends and in the main text. Reported n values can be found in the figure legends and in the results. All data are represented as mean ± s.e.m. (standard error of mean) unless indicated otherwise. All replicates were biological. Sample sizes were not *a priori* determined and are in line with (or exceeding) standards in the field. All statistical tests are indicated in the respective figure legend and are two-sided. In all figure legends the exact value and definition of n is indicated. In all panels: not significant, ns p > 0.05; * p < 0.05; ** p < 0.01; *** p < 0.001.

Differential gene expression tests were performed with exact test for negative binomial distribution implanted in the R package edgeR, and results were filtered for log2FC (fold change) and p-value: log2FC > 1 and p-value < 0.05.

#### Gene Ontology (GO)

Gene Ontology (GO) annotations (Ashburner et al., 2000; Gene Ontology, 2021) were determined using the R package limma (v.3.44.3) and custom R scripts. GO terms were taken from the Bioconductor annotation package org.Mm.eg.db (v.3.11.4) released on 2020-05-02. The following GO terms were selected: detection of mechanical stimulus (GO:0050982), ion channel activity (GO:0005216) and integral component of membrane (GO:0016021).

## Materials availability

This study did not generated new unique reagents.

## Data and code availability

The RNA-seq datasets generated and analyzed have been deposited in NCBI’s Gene Expression Omnibus (Edgar et al., 2002) and are accessible through the following GEO Series accession numbers: GSE168032.

## Notes

### Competing Interest Statement

The authors have declared no competing interest.

