## Supplemental information for "Patch-seq of mouse DRG neurons reveals candidate genes for specific mechanosensory functions"

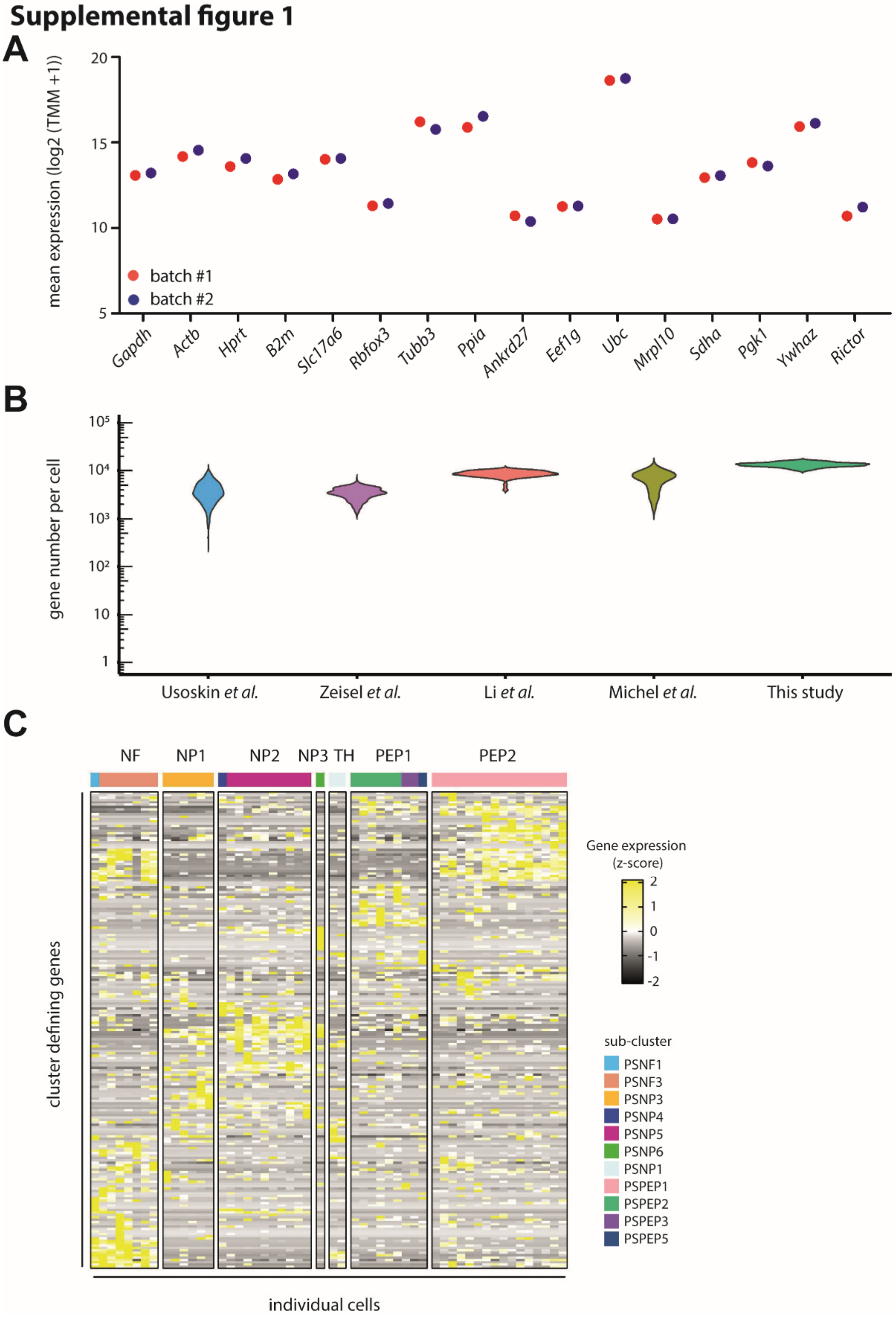


**Supplemental Figure 1. RNA Sequencing Data Quality**

(**A**) Expression levels of representative reference genes of the two sequencing batches. Data shown are means of log2 transformed TMM+1 values. (**B**) Violin plot of number of expressed genes per DRG neuron in DRG RNA sequencing studies, as indicated. Expressed genes are set with a threshold of TPM ≥ 1 except for Usoskin dataset for which the threshold is CPM ≥ 1. (**C**) Heat map of expression of the 211 genes used for DRG neuron clustering (see **Fig.2A**).

Supplemental figure 2

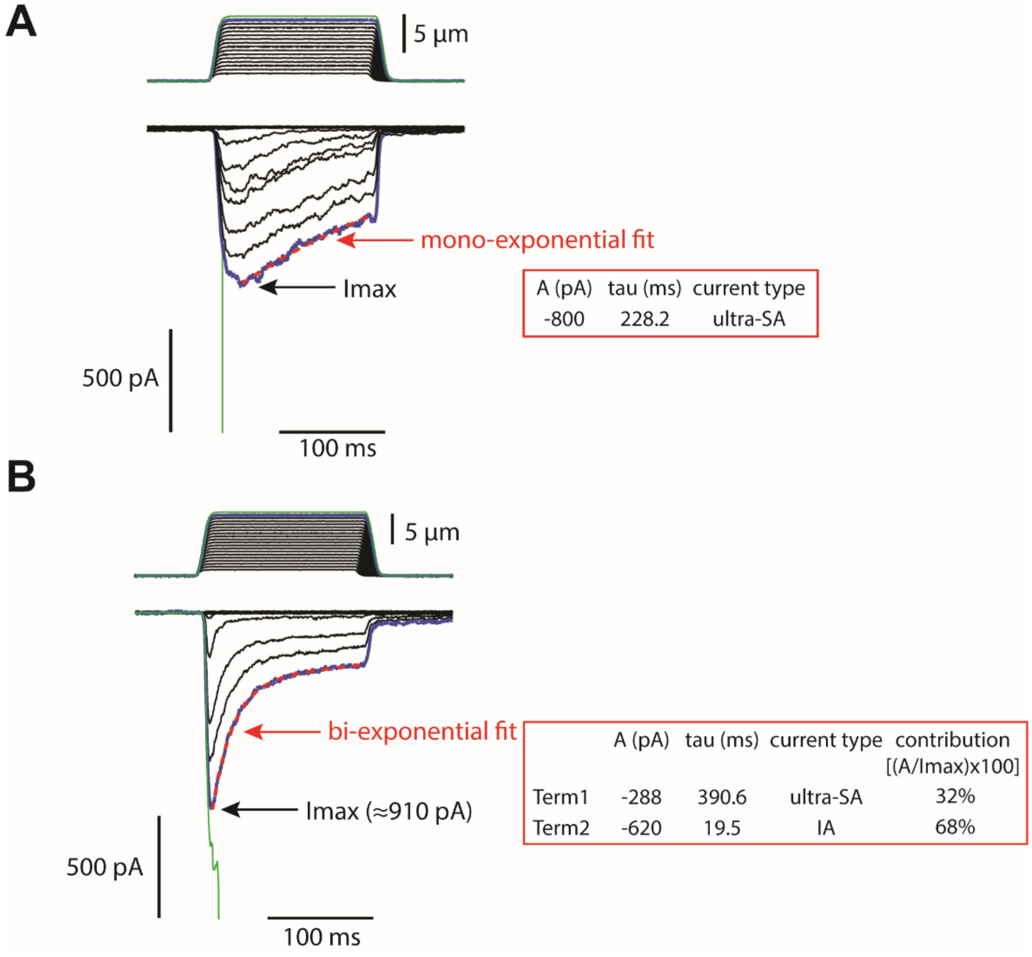


**Supplemental Figure 2. Analysis of mono- and bi-phasic MS currents**

Typical current traces of monophasic (**A**) and biphasic (**B**) MS currents. Currents were elicited by increasing mechanical stimulus up to patch rupture (green traces). Inactivation kinetics were fitted with mono- (**A**) or bi-exponential (**B**) equation (red dashed lines), giving fitting parameters as depicted. Tau values were used to classify MS current types. For biphasic currents, the contribution of each MS current type is determined from the ratio A/Imax (*100).

### Supplemental figure 3

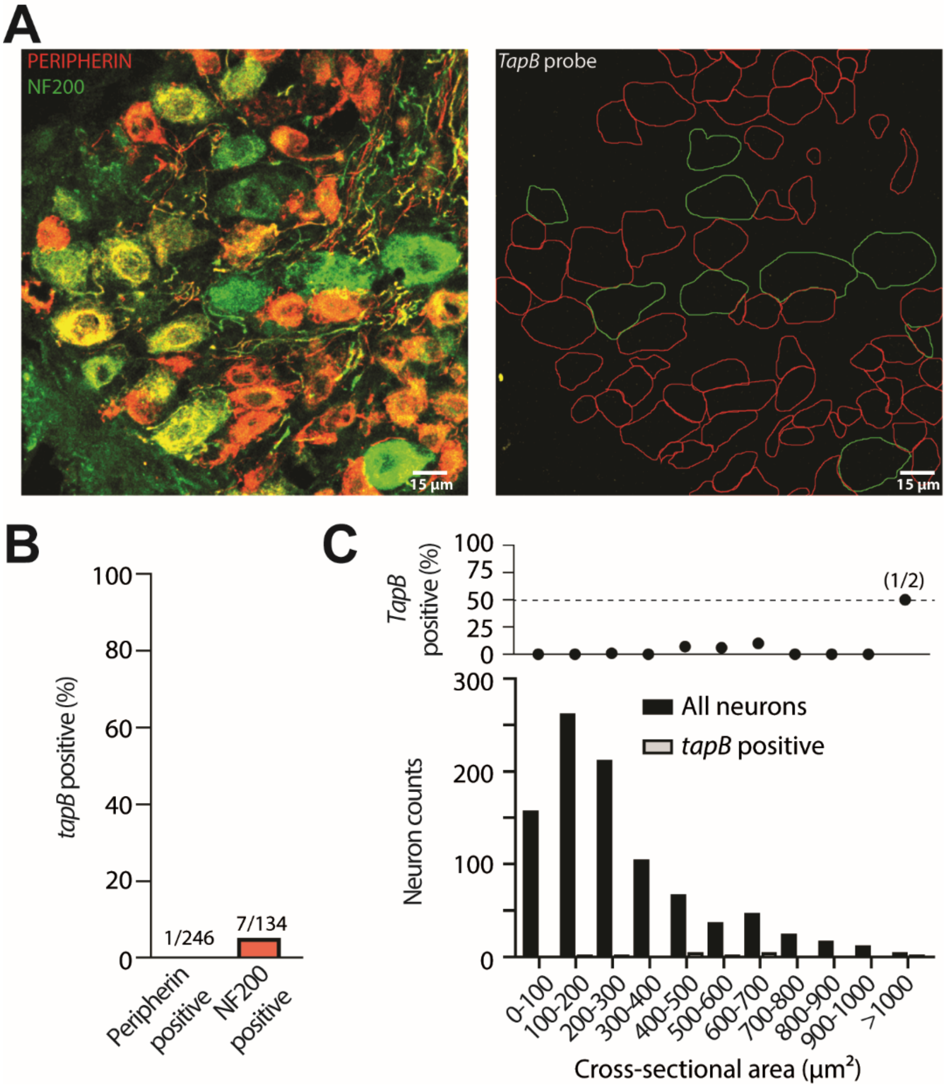


**Supplemental Figure 3.** ***In Situ*** **Hybridization negative control, related to Figure 5A-C**

(**A**) Representative images of fluorescent *in situ* hybridization for the control gene *TapB* (right panel) in DRG neurons immuno-stained for peripherin and NF200 (left panel). (**B**) Percentage of *ISH* positive DRG neurons in Peripherin- and NF200- positive populations. (**C**) Cross-sectional area distribution of *TapB* mRNA positive neurons. Top panel shows the percentage of *TapB* positive neurons.

### Supplemental figure 4

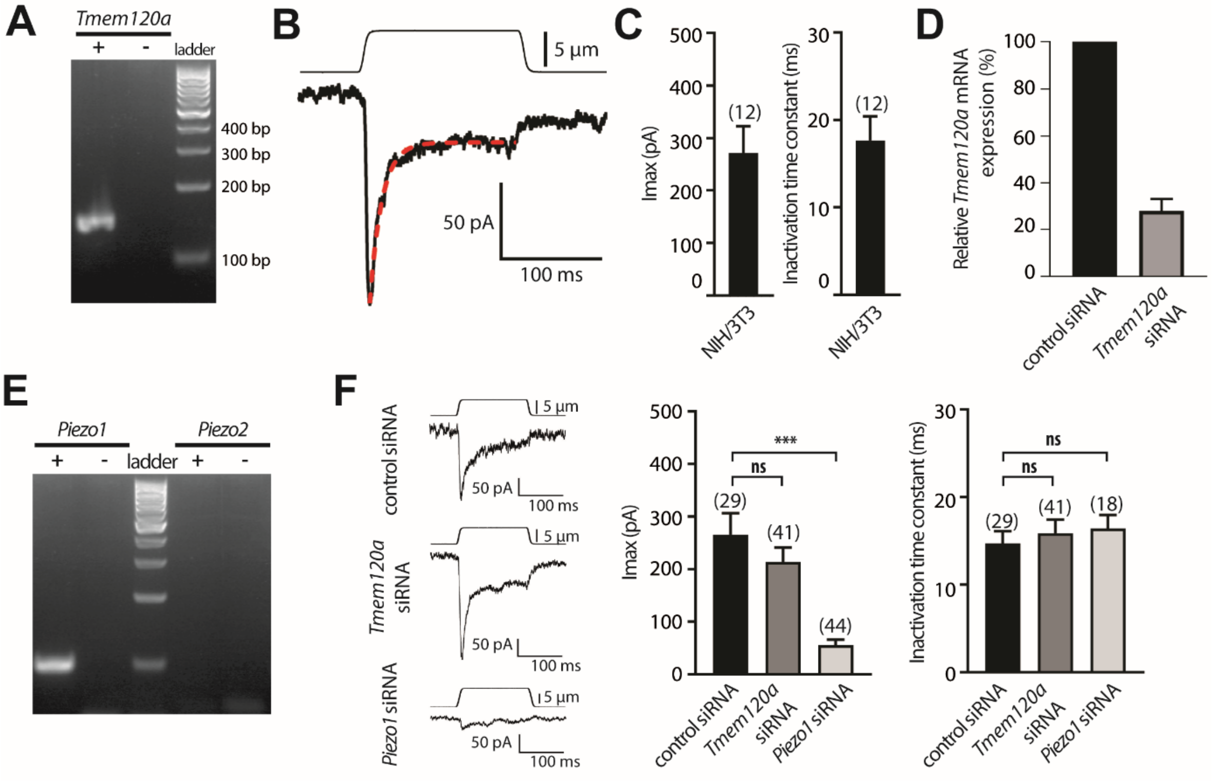


**Supplemental Figure 4. Efficient downregulation of *Tmem120a* has no effect on endogenous MS current in NIH/3T3 cells, related to Figure 5D-G**

(**A**) PCR products in NIH/3T3 cells using *Tmem120a* specific primers from reverse transcription done in the presence (+) or absence (–) of RT enzyme. (**B**) Representative recording of a MS current elicited at a holding potential of -80 mV in NIH/3T3 cells. Red dashed line represents fit of inactivation with a mono-exponential equation ($\tau$ = 11.7 ms). (**C**) Average maximal amplitude (left panel) and time-constant of inactivation (right panel) of NIH/3T3 MS current elicited at holding potential of -80 mV. (**D**) Expression of *Tmem120a* mRNA in NIH/3T3 cells determined by RT-qPCR 48h after electroporation with control or *Tmem120a* siRNA (n = 2, *Gapdh* and *β-actin* as reference genes). Electroporated and unelectroporated cells are not sorted leading to underestimation of down-regulation. (**E**) PCR products in NIH/3T3 cells using *Piezo1* and *Piezo2* specific primers from reverse transcription performed in the presence (+) or absence (–) of RT enzyme. (**F**) Typical recording traces (left panels), average maximal amplitude (middle panel) and time-constant of inactivation (right panel) of MS currents elicited at holding potential of -80 mV in NIH/3T3 cells transfected with control, *Tmem120a* or *Piezo1* siRNA. In panel B and F, upper traces represent the mechanical probe displacement and lower traces the recorded currents. N numbers are indicated in brackets. Error bars represent s.e.m.; ***, $p < 0.001$; ns, not significant; Kruskal-Wallis multiple comparison test.

### Supplementary figure 5

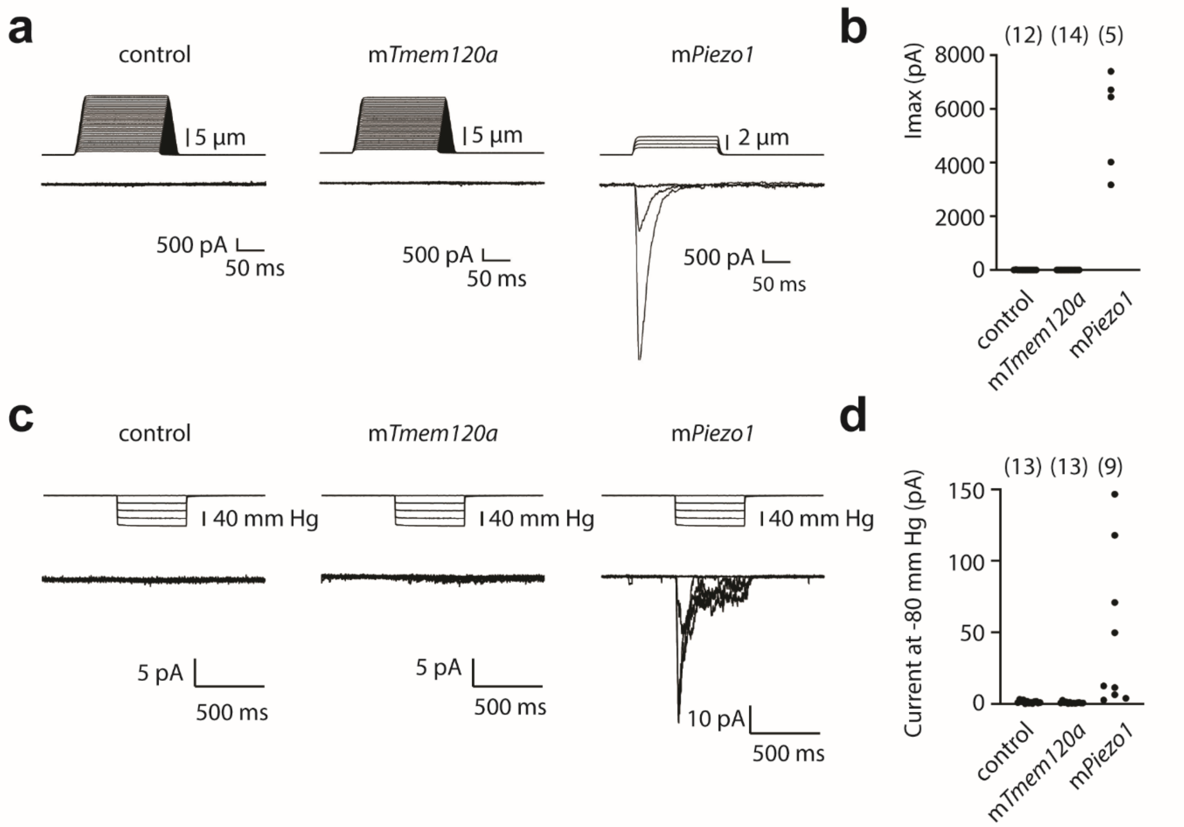


**Supplemental Figure 5. Overexpression of mouse *Tmem120a* does not induce MS current in HEK-P1KO cells**

**(A)** Representative examples of recordings in cells voltage clamped at -80 mV and stimulated with a mechanical probe under whole-cell configuration. Cells were transiently transfected with control vector, *Tmem120a* or *Piezo1*, as specified. (**B**) Maximal MS current amplitude in each stimulated cell. N numbers are in brackets. **(C)** Representative examples of recordings in cell-attached patches stimulated with pulses of negative pressure applied in the recording pipette. Cells were transiently transfected with control vector, *Tmem120a* or *Piezo1*, as specified. (**D**) Maximal MS current amplitude elicited with -80 mm Hg stimulation step in each stimulated cell. N numbers are in brackets.

| Cell number | Neuron type | Genetic cluster (Nomenclature from Zeisel *et al.*, 2018) | Mixed currents | Current type 1 (major) | Current type 2 (minor) | Proportions [type1 (%) / type2 (%)] |
|---|---|---|---|---|---|---|
| 1 | **NF** | PSNF1 | NO | RA | | |
| 2 | **NF** | PSNF3 | NO | RA | | |
| 3 | **NF** | PSNF3 | NO | RA | | |
| 4 | **NF** | PSNF3 | NO | RA | | |
| 5 | **NF** | PSNF3 | NO | RA | | |
| 6 | **NF** | PSNF3 | NO | RA | | |
| 7 | **NF** | PSNF3 | NO | RA | | |
| 8 | **NF** | PSNF3 | YES | RA | US | 78/22 |
| 9 | **NP1** | PSNP3 | YES | IA | US | 69/31 |
| 10 | **NP1** | PSNP3 | YES | IA | US | 82/18 |
| 11 | **NP1** | PSNP3 | YES | RA | US | 62/38 |
| 12 | **NP1** | PSNP3 | YES | US | IA | 54/46 |
| 13 | **NP1** | PSNP3 | NO | SA | | |
| 14 | **NP1** | PSNP3 | YES | SA | US | 72/28 |
| 15 | **NP2** | PSNP4 | NO | RA | | |
| 16 | **NP2** | PSNP5 | NO | RA | | |
| 17 | **NP2** | PSNP5 | YES | RA | SA | 88/12 |
| 18 | **NP2** | PSNP5 | YES | RA | SA | 86/14 |
| 19 | **NP2** | PSNP5 | YES | RA | SA | 88/12 |
| 20 | **NP2** | PSNP5 | NO | IA | | |
| 21 | **NP2** | PSNP5 | NO | IA | | |
| 22 | **NP2** | PSNP5 | YES | IA | US | 75/25 |
| 23 | **NP2** | PSNP5 | YES | US | IA | 59/41 |
| 24 | **NP2** | PSNP5 | YES | IA | US | 67/33 |
| 25 | **NP2** | PSNP5 | NO | US | | |
| 26 | **NP3** | PSNP6 | YES | US | IA | 82/18 |
| 27 | **TH** | PSNP1 | NO | RA | | |
| 28 | **TH** | PSNP1 | NO | US | | |
| 29 | **PEP1** | PSPEP2 | NO | IA | | |
| 30 | **PEP1** | PSPEP2 | YES | IA | US | 90/10 |
| 31 | **PEP1** | PSPEP2 | YES | IA | US | 78/22 |
| 32 | **PEP1** | PSPEP2 | YES | IA | US | 68/32 |
| 33 | **PEP1** | PSPEP2 | | NR | | |
| 34 | **PEP1** | PSPEP2 | | NR | | |
| 35 | **PEP1** | PSPEP3 | NO | RA | | |
| 36 | **PEP1** | PSPEP3 | NO | RA | | |
| 37 | **PEP1** | PSPEP5 | | NR | | |
| 38 | **PEP2** | PSPEP1 | NO | RA | | |
| 39 | **PEP2** | PSPEP1 | YES | RA | US | 67/33 |
| 40 | **PEP2** | PSPEP1 | YES | RA | SA | 58/42 |
| 41 | **PEP2** | PSPEP1 | NO | RA | | |
| 42 | **PEP2** | PSPEP1 | YES | RA | US | 73/27 |
| 43 | **PEP2** | PSPEP1 | YES | IA | US | 72/28 |
| 44 | **PEP2** | PSPEP1 | NO | SA | | |
| 45 | **PEP2** | PSPEP1 | YES | SA | RA | 69/31 |
| 46 | **PEP2** | PSPEP1 | YES | US | RA | 75/25 |
| 47 | **PEP2** | PSPEP1 | NO | US | | |
| 48 | **PEP2** | PSPEP1 | NO | SA | | |
| 49 | **PEP2** | PSPEP1 | YES | US | RA | 67/33 |
| 50 | **PEP2** | PSPEP1 | NO | US | | |
| 51 | **PEP2** | PSPEP1 | YES | SA | RA | 51/49 |
| 52 | **PEP2** | PSPEP1 | NO | US | | |
| 53 | **PEP2** | PSPEP1 | NO | US | | |

**Supplemental Table 1. Summary of transcriptomics classification and mechanoelectrical properties of the scRNAseq neurons**

RNA sequenced DRG neurons are labeled according to their genetically identified neuronal type and their MS currents characterized by patch clamp experiments. Mixed currents are MS currents for which inactivation kinetics could be fitted with bi-exponential function, revealing the presence of two distinct MS components, with the smaller (current type 2) contributing to at least 10% of the peak current amplitude.

**Supplemental Table 2. Set of transcripts enriched in neurons grouped by MS current types**
